# Quiescent cell re-entry is limited by macroautophagy-induced lysosomal damage

**DOI:** 10.1101/2024.10.11.617892

**Authors:** Andrew Murley, Ann Catherine Popovici, Xiwen Sophie Hu, Anina Lund, Kevin Wickham, Jenni Durieux, Larry Joe, Etai Koronyo, Andrew Dillin

## Abstract

To maintain tissue homeostasis, many cells are kept in a quiescent state until prompted to divide. The re-activation of quiescent cells is perturbed with aging and may underlie declining tissue homeostasis and resiliency. The unfolded protein response regulators IRE-1 and XBP-1 are required for the re-activation of quiescent cells in developmentally L1 arrested *C. elegans.* Utilizing a forward genetic screen in *C. elegans*, we discovered that macroautophagy targets protein aggregates to lysosomes in quiescent cells, leading to lysosome damage that prevents cell cycle re-entry in the absence of IRE-1/XBP-1. Genetic inhibition of macroautophagy and stimulation of lysosomes via the overexpression of HLH-30 (TFEB/TFE3) synergistically reduces lysosome damage. Protein aggregates are also targeted to lysosomes by macroautophagy in quiescent cultured mammalian cells, causing lysosome damage that is associated with reduced re-activation. Thus, lysosome damage is a hallmark of quiescent cells and limiting lysosome damage by restraining macroautophagy can stimulate their re-activation.

## Introduction

Age is associated with a decline in tissue function and the diminished capacity for repair ^1^. Adult somatic stem cells, such as hematopoietic stem cells, neural stem cells, and muscle stem cells give rise to new cell types within blood, brain and muscle tissues, respectively ^2,3^. These adult stem cells divide infrequently, mainly residing in a quiescent (G_0_) cell state until called upon to proliferate in response to specific conditions^3^. Quiescence is a general regulator of tissue and organismal homeostasis, and is an important feature of other cells in the body, including lymphocytes and hepatocytes ^2^. Studies in model organisms and humans indicate that aging, and age-associated diseases, compromise the appropriate re-activation of quiescent cells, which may contribute to progressive tissue dysfunction with age ^4–8^. The mechanisms leading to compromised adult stem cell function with age are incompletely understood but are influenced by cell intrinsic properties and signals from their niche ^3,4^. Identifying processes that contribute to the age-associated decline in stem cell function may identify ways to harness the body’s innate capacity for repair and rejuvenation throughout life.

Cellular quiescence is an ancient homeostatic mechanism eukaryotic cells employ in response to conditions that are not conducive to growth either to maintain tissue homeostasis or because essential nutrients are limiting. In single-celled eukaryotes such as yeast and in simpler animals such as *C. elegans*, one of the main signals inducing cellular quiescence is nutrient deprivation. When born in the absence of food, *C. elegans* L1 larvae arrest their development until they encounter sufficient food sources, termed L1 arrest. At the cellular level, during L1 arrest, reduced Insulin-like growth factor (IGF) signaling activates the transcription factor DAF-16 (FOXO), inducing transcriptional upregulation of the cyclin-dependent kinase inhibitor CKI-1/2 (p21/p27), which arrests the post-embryonic division of cells in the animal ^9–11^. *C. elegans* can survive L1 arrest for a few weeks, however, over time animals lose the ability to develop into adults, remaining permanently arrested as L1 larvae^12^. Cellular integrity plays a key role in the ability of animals to recover from prolonged L1 arrest, with animals that accumulate more reactive-oxygen species (ROS), protein aggregates and fragmented mitochondria over time being unable to re-enter development ^12^. These signs of cellular dysfunction resemble those found in aging cells, suggesting that the mechanisms that lead to permanent arrest of *C. elegans* L1 larvae may be similar to those restricting the reactivation of quiescent cells in aged mammals, such as adult somatic stem cells. Being a multicellular model for studying quiescence, *C. elegans* offers the opportunity to explore how intercellular signaling influences homeostatic mechanisms and re-activation in quiescent cells, which is an advantage over homogenous cell culture models.

Numerous homeostatic mechanisms maintain the function of quiescent cells. Being mitotically arrested, quiescent cells generally have lower anabolic rates, which is maintained in part by reduced mechanistic target-of-rapamycin (mTOR) activity ^13–15^. Reduced mTOR signaling in quiescent cells helps to inhibit their growth and transition to cellular senescence, which is characterized by high mTOR signaling that is resistant to changes in nutrient levels ^16^. Aging compromises the autophagy/lysosomal pathways, and various means of counteracting this decline, mainly through stimulation of the transcription factor Tfeb, improve quiescent cell homeostasis and re-activation efficiency ^5–7^. The unfolded protein response of the endoplasmic reticulum (UPR) also plays a role in quiescent cell homeostasis. The UPR promotes homeostasis of the ER and responds to increased cellular need for the ER by activating the IRE-1/XBP-1, ATF6 and PERK pathways, reviewed extensively elsewhere ^17–21^. Activation of the UPR is diminished in certain cells and tissues during aging, and previous studies have reported that IRE-1 is required for the development of *C. elegans* after prolonged periods of L1 arrest and that Ire1α is crucial for liver regenerative responses in mice, indicating that diminished quiescent cell re-activation with aging might be influenced by aberrant UPR signaling ^12,22^..

In this study, we report that damage to lysosomes is a conserved feature of quiescent cells and compromises their re-activation in the absence of IRE-1 or XBP-1. Macroautophagy makes an important contribution to this damage through the targeting of protein aggregates to lysosomes, and conversely, boosting lysosome biogenesis and inhibiting macroautophagy work synergistically to improve the re-activation of quiescent cells in *C. elegans* and mammalian cells. Because quiescent cell re-activation declines with age, our results suggest that, rather than boosting macroautophagy, one strategy to combat this age-associated decline could be to boost lysosome function while restraining macroautophagy.

## Results

### Loss-of-function mutations in core macroautophagy genes improves recovery of animals from L1 arrest

Consistent with previously published findings, *C. elegans* can survive L1 arrest for a few weeks, but over time lose the ability to develop once fed. This phenotype is accelerated in *ire-1* and *xbp-1* mutants, encoding two main transducers of the UPR (**Figure 1A-B**) ^12^. Consistent with previous observations, *atf-6* and *pek-1* mutations did not affect recovery from L1 arrest (**Figure 1B**) ^12^. The phenotype of *ire-1* and *xbp-1* mutants was also distinct from *hlh-30* mutants, encoding the *C. elegans* ortholog of TFEB, which died rapidly during L1 arrest (**Figure 1B**). The ability of *ire-1* and *xbp-1* mutants to survive quiescence, but over time lose the ability to escape it, is similar to observations made in a variety of adult stem cells and other quiescent cells in vertebrates whose ability to escape quiescence declines with aging and disease ^4–6,8,23^. Notably, Ire1α is also required for re-activation of quiescent hepatocytes in damaged livers of mice ^22^. This phenotype, which is present in WT animals, but exacerbated in *ire-1* and *xbp-1* mutants, we term “progressive terminal L1 arrest” or PTLA. The inability of *ire-1* mutants to recover from PTLA is correlated with persistent expression of the cell-cycle inhibitor CKI-1 in seam cells, a cell type that becomes arrested during L1 arrest, indicating that persistent UPR dysfunction during cellular quiescence prevents replication-competent cells in the animals from escaping quiescence and re-entering the cell cycle (**Figure 1C**) ^10^.

**Figure 1.**
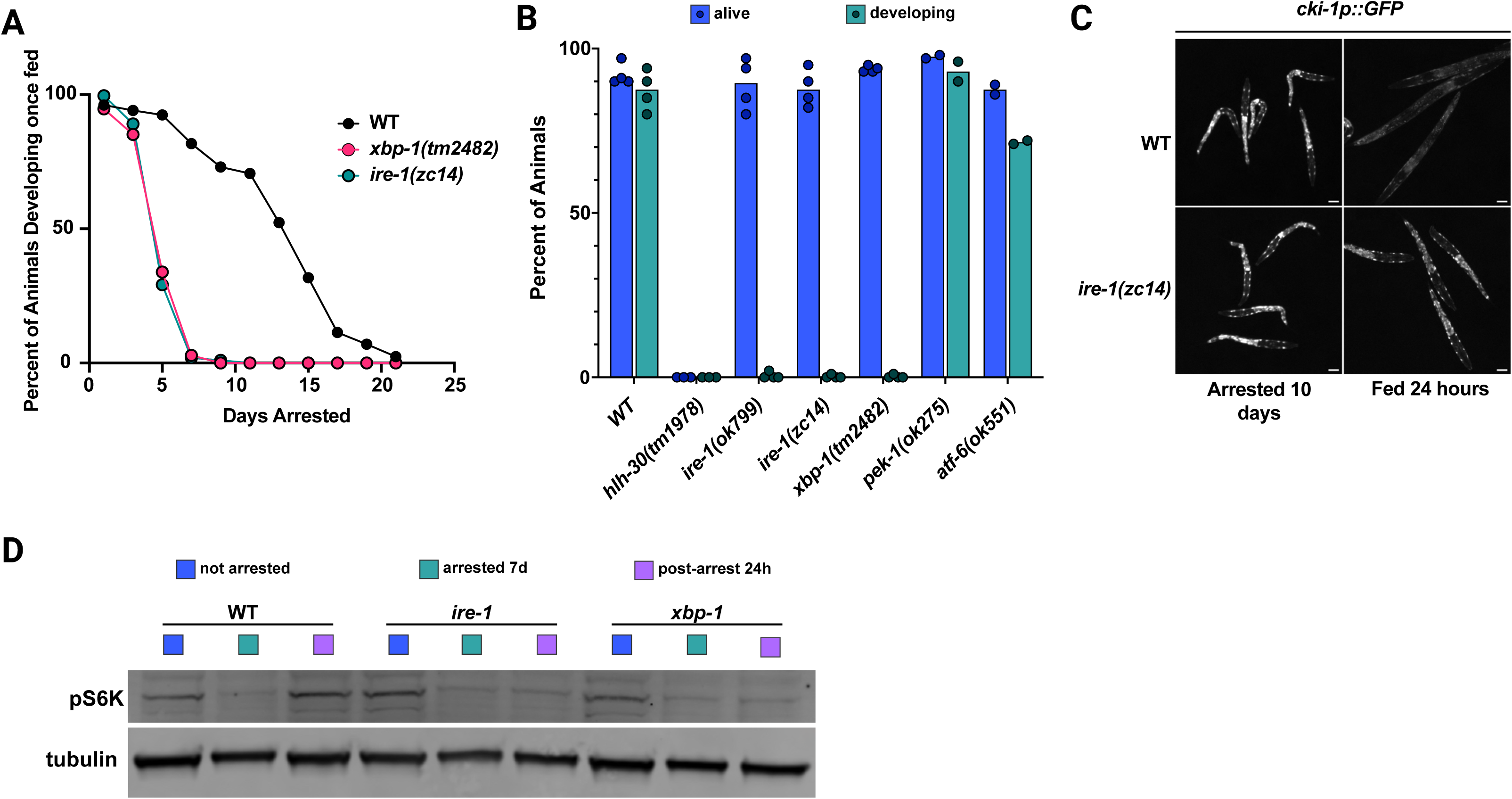
– IRE-1 and XBP-1 are required for the re-activation of mTORC1 after prolonged L1 arrest. **(A)** WT, *ire-1(zc14)*, and *xbp-1(tm2482)* mutants were subjected to L1 arrest, and every other day, animals were given food and scored 2-4 days later for their ability to develop to the L3 larval stage. **(B)** Animals of the indicated genotypes were subjected to L1 arrest for 10 days, then given food and assessed for their ability to develop to the L3 larval stage after 2-4 days. **(C)** WT and *ire-1* animals expressing GFP driven by the CKI-1 promoter (*cki-1p::GFP)* were subjected to L1 arrest for 10 days and then given food and imaged. Scale bars = 20 μm. **(D)** Representative Western blot of pS6K and α-tubulin. Fed L1 animals (not arrested), animals L1 arrested for 7 days (arrested 7d), and animals L1 arrested for 7 days then fed for 24 hours (post-arrest 24h) were harvested as described in “Materials and Methods” and subjected to Western Blot analysis.

Interestingly, despite being able to ingest food after prolonged L1 arrest, *ire-1* and *xbp-1* mutants cannot re-activate mechanistic target of rapamycin 1 (mTORC1) signaling as assessed by phosphorylation of RSKS-1, the *C. elegans* S6 kinase (**Figure 1D and Figure S1A**) ^24,25^. As a major sensor of nutrients and growth factors, mTORC1 is a kinase that is active in conditions permissive to cell growth, phosphorylating and activating proteins promoting cell growth, including those involved in protein translation and ribosome biogenesis, and inhibiting catabolic processes such as autophagy ^26^. Thus, aberrant regulation of mTORC1 in *ire-1* and *xbp-1* mutant animals and persistent expression of cell cycle inhibitors in arrested cells were correlated with their inability to develop after prolonged L1 arrest.

To better understand the mechanisms leading to progressive development of a permanent cell cycle arrest in L1-arrested *ire-1* mutants, we conducted a forward genetic screen combined with whole genome sequencing to identify mutations that would bypass the requirement for IRE-1 in L1 arrest. We isolated F2 progeny from EMS-mutagenized *ire-1(ok799)* and *ire-1(zc14)* animals and subjected them to L1 arrest for a period during which all *ire-1* mutants lose the ability to develop but WT animals remain capable of developing. After L1 arrest, we allowed animals to recover on plates with food. Animals that reached adulthood and were fertile were used to establish lines that were re-tested to ensure reproducibility, then subjected to whole-genome sequencing to identify candidate mutations (**Figure 2A**).

**Figure 2.**
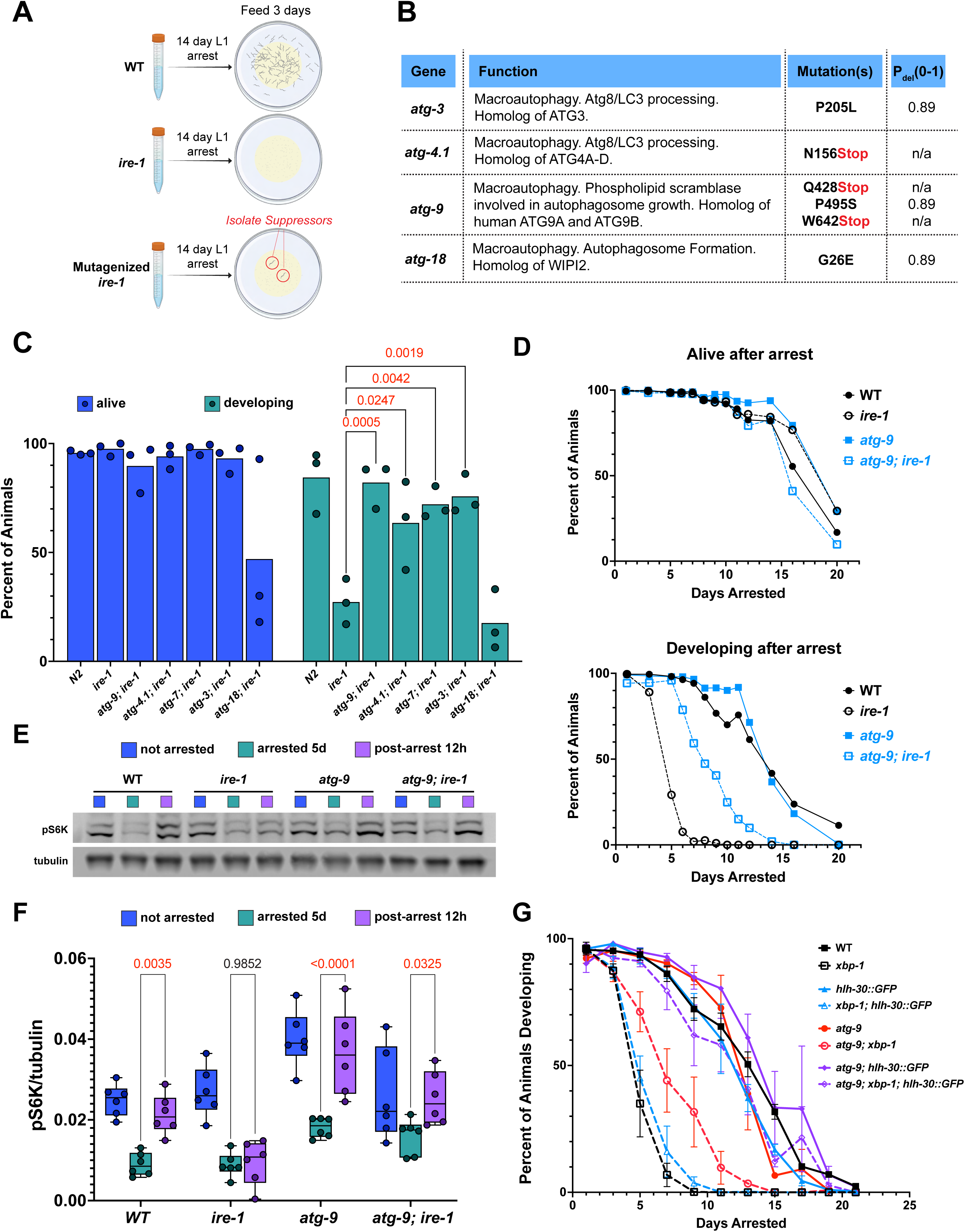
– Mutations in core macroautophagy genes promotes recovery from L1 arrest and restores mTORC1 regulation. **(A)** Overview of forward genetic screen for suppressors of PTLA phenotype in *ire-1* mutants, where F2 progeny from EMS-mutagenized *ire-1* mutants for L1 arrested for 14 days, then plated on plates with food. Rare mutants that developed to adulthood and were fertile were used to establish lines that were analyzed by whole-genome sequencing. **(B)** Mutations isolated from the screen. “P_del_” represents the probability the indicated mutation is deleterious to the encoded protein based on PANTHER protein database. **(C)** Loss-of-function mutations in macroautophagy genes suppresses PTLA. *ire-1* mutants were crossed to *atg-3(bp412), atg-4.1(bp501), atg-7(bp422), atg-9(bp564)* and *atg-18(gk378)* and subjected to a five-day L1 arrest. The percentage of animals surviving L1 arrest and developing past L1 larval stage after 4 days of feeding were assessed from 150-200 animals from three independent biological replicates. P values calculated using Dunnett’s multiple comparisons test. **(D)** *atg-9* mutation suppresses PTLA. Animals of the indicated genotypes were subjected to L1 arrest and periodically assessed for survival and ability to develop to the L3-L4 larval stage **(E)** Representative Western blot of pS6K and α-tubulin. Fed L1 animals (not arrested), animals L1 arrested for 5 days (arrested 5d), and animals L1 arrested for 5 days then re-fed for 12 hours (post-arrest 12h) were harvested as described in “Methods” and subjected to Western Blot analysis. **(F)** Quantification of pS6K:tubulin ratios in animals as in (D). Bar graphs represent mean values overlaid with individual values. P values (in red) were calculated from n = 6 independent biological replicates using Tukey’s multiple comparison test from a two-way ANOVA. **(G)** Reduced autophagy and HLH-30∷GFP overexpression synergistically suppress PTLA. Animals of the indicated genotypes were subjected to L1 arrest and periodically assessed for survival and ability to develop to the L3 larval stage. Error bars represent standard error of the mean (SEM) for n=3 biological replicates.

Amongst the genes identified in our screen were, surprisingly, strong loss-of-function alleles in macroautophagy genes (**Figure 2B**). Macroautophagy (herein “autophagy”) is an ancient mechanism eukaryotic cells employ to engulf superfluous, damaged, or toxic cellular components in double-membrane structures called autophagosomes, which fuse with lysosomes that degrade autophagosome cargo ^27^. We isolated lines with three independent mutations in *atg-9*, two resulting in pre-mature stop codons (*atg-9(Q428Stop)* and *atg-9(W642Stop)*) and one in a highly conserved proline residue that, based on structural homology, likely generates a loss of function allele (*atg-9(P495S))* (**Figure 2B**). ATG-9 is the sole transmembrane protein required in the core autophagic machinery and is localized to vesicles that are recruited to the phagophore assembly site (PAS) and serve a crucial role in initiating autophagy. Biochemical and structural studies indicate that yeast and human Atg9 form a homotrimeric complex that functions as a phospholipid scramblase, and *atg-9* is the sole homolog of yeast *ATG9* in the *C. elegans* genome ^28–31^. We also isolated a premature stop codon in one of two *C. elegans* homologs of yeast *ATG4*, *atg-4.1* (*atg-4.1(N156Stop)*) (**Figure 2B**). Akin to yeast Atg4, ATG-4.1 is a cysteine peptidase that is required for processing of LGG-1 (Atg8/LC3), the core component of autophagic membranes. In *C. elegans,* ATG-4.1 and its paralog ATG-4.2 have somewhat overlapping functions in autophagy, and double *atg-4.1;atg-4.2* mutants are not viable, but each are more important for autophagy of certain substrates in specific cell types or conditions and ATG-4.1 has much higher enzymatic activity than ATG-4.2 ^32,33^. Other mutations we identified in *atg-*3, involved in lipidation of LGG-1, and *atg-18* are predicted to be detrimental to the function of the proteins encoded by those genes (**Figure 2B**). Notably, all the proteins encoded by these genes function early in the autophagy process, *i.e.,* in autophagosome formation, but not in lysosomal fusion of autophagosomes or lysosomal degradation processes.

We were surprised to identify such strong hits in core autophagy machinery in our screen, since L1 arrest is a starvation response that is thought to require autophagy for organismal survival ^34^. To confirm the results from our screen, we crossed *ire-1* animals to independently isolated lines harboring null mutations in *atg-9*, *atg-4.1* and *atg-18,* as well as hypomorphic alleles of the essential genes *atg-3* and *atg-7*, which regulate distinct steps in autophagosome formation ^35^. Although a null mutation in *atg-18* compromised survival of L1-arrested animals, other mutations compromising autophagy improved the recovery of L1-arrested *ire-1* mutants after five days of arrest, strongly suggesting that autophagy, and no other cellular functions of autophagy proteins is relevant to PTLA (**Figure 2C**) ^36^. Although mutations in autophagy improved PTLA in *ire-1* mutants, it did not fully restore *ire-1* mutants to a WT PTLA phenotype, indicating that other processes also influence PTLA (**Figure 2D**).

Mitochondrial DNA (mtDNA) is degraded by autophagy during L1 arrest, and subsequent mtDNA synthesis and mitochondrial biogenesis via the mitochondrial UPR influences recovery from L1 arrest ^37,38^. Given the role the ER plays in mtDNA replication and segregation in cells, IRE-1 and XBP-1 may play a role in mtDNA replication, and therefore, blocking autophagy may be beneficial by preventing the degradation of mtDNA ^39,40^. To test this idea, we asked if mutations in *pdr-1*, the *C. elegans* ortholog of *Parkin* involved specifically in autophagy of mitochondria (mitophagy), could rescue PTLA in *ire-1* mutants ^41^. We found that *pdr-1* mutants had no effect on PTLA in *ire-1* mutants (**Figure S2A**).

To assess whether blocking autophagy corrects mTORC1 regulation in *ire-1* mutants, we analyzed phosphorylation of RSKS-1 in un-arrested L1 animals, L1 animals arrested five days, and L1 animals arrested for 5 days and fed for 12 hours. Consistent with their improved development after L1 arrest, *atg-9; ire-1* mutants also had restored activation of mTORC1 following re-feeding (**Figure 2E-F**). Relative levels of RSKS-1 phosphorylation during arrest also appeared slightly higher in *atg-9* mutants but were not statistically significant (**Figure 2F**). Consistent with our finding is the role of autophagy in regulating the activation of mTORC1 in response to T-cell receptor stimulation in Treg cells ^42^.

### Autophagy inhibition and lysosome biogenesis synergistically improve recovery from L1 arrest

Autophagy is a tightly regulated process in cells, and excessive autophagy can lead to cell death or possibly to the degradation of cellular constituents necessary for cell cycle re-entry ^43^. To test if excessive autophagy is detrimental to PTLA in *ire-1 or xbp-1* mutants, we overexpressed the *C. elegans* ortholog of *Tfe3/Tfeb* transcription factors, HLH-30, fused to GFP. This family of transcription factors boost the overall autophagic capacity of cells by upregulating genes for lysosome function and autophagosome formation, and consistent with this, overexpression of HLH-30 stimulates autophagic flux in *C. elegans* ^44–46^. We found that overexpression of HLH-30::GFP had a marginal, slightly positive effect on PTLA in *xbp-1* mutants (**Figure 2G**).

As HLH-30 overexpression boosts autophagic capacity of cells via increased lysosome biogenesis/function and autophagosome formation, we tested if the increased flux through the autophagy pathway prior to lysosomal fusion is compensated for by an increase in lysosome function and if stimulating lysosome function while restraining autophagy may have added beneficial effects. We found that while *xbp-1* animals overexpressing HLH-30::GFP or harboring a null mutation in *atg-9* were somewhat resistant to PTLA, *xbp-1* animals combining both interventions were remarkably resistant to PTLA, developing after prolonged periods of L1 arrest almost as well as WT animals (**Figure 2G**). WT animals combining both interventions were also slightly more resistant to PTLA after 9-13 days of L1 arrest (**Figure 2G**).

### Autophagy leads to lysosome damage in L1-arrested animals

These findings suggested to us that autophagy during quiescence may compromise lysosomal function, possibly by delivering difficult to degrade substances such as protein aggregates and may underly dysfunctional mTORC1 re-activation and re-entry into the cell cycle. The consequences of lysosomal protein aggregates can be counteracted by increasing lysosomal function and biogenesis via increased HLH-30 expression, albeit with the counterproductive downside of increased autophagosome formation as well. Consistent with this idea, reducing autophagy of protein aggregates in *sqst-1* and, to a lesser extent, *epg-7* mutants mitigated PTLA (**Figure S2B-C**) ^47^. Furthermore, quiescent neural stem cells harboring intra-lysosomal protein aggregates have poor re-activation efficiency that is boosted by various interventions that promote clearance of lysosomal protein aggregates ^5^. Additionally, damaging lysosomes in mammalian cells has been shown to inactivate mTORC1 via altered regulation of the Rag-Ragulator pathway ^48^.

We used proteostat dye, which specifically binds to misfolded and aggregated proteins, and a lysosomal membrane marker, SCAV-3::GFP, to test if protein aggregates are associated with lysosomes in L1 arrested animals ^49^. We found that protein aggregates were commonly associated with lysosomes in WT animals during L1 arrest (**Figure 3A-B, D**). In contrast, the association of protein aggregates with lysosomes was greatly diminished in *atg-9* mutants (**Figure 3A-B**). Lysosomes were also smaller in *atg-9* animals compared to WT (**Figure 3C**).

**Figure 3.**
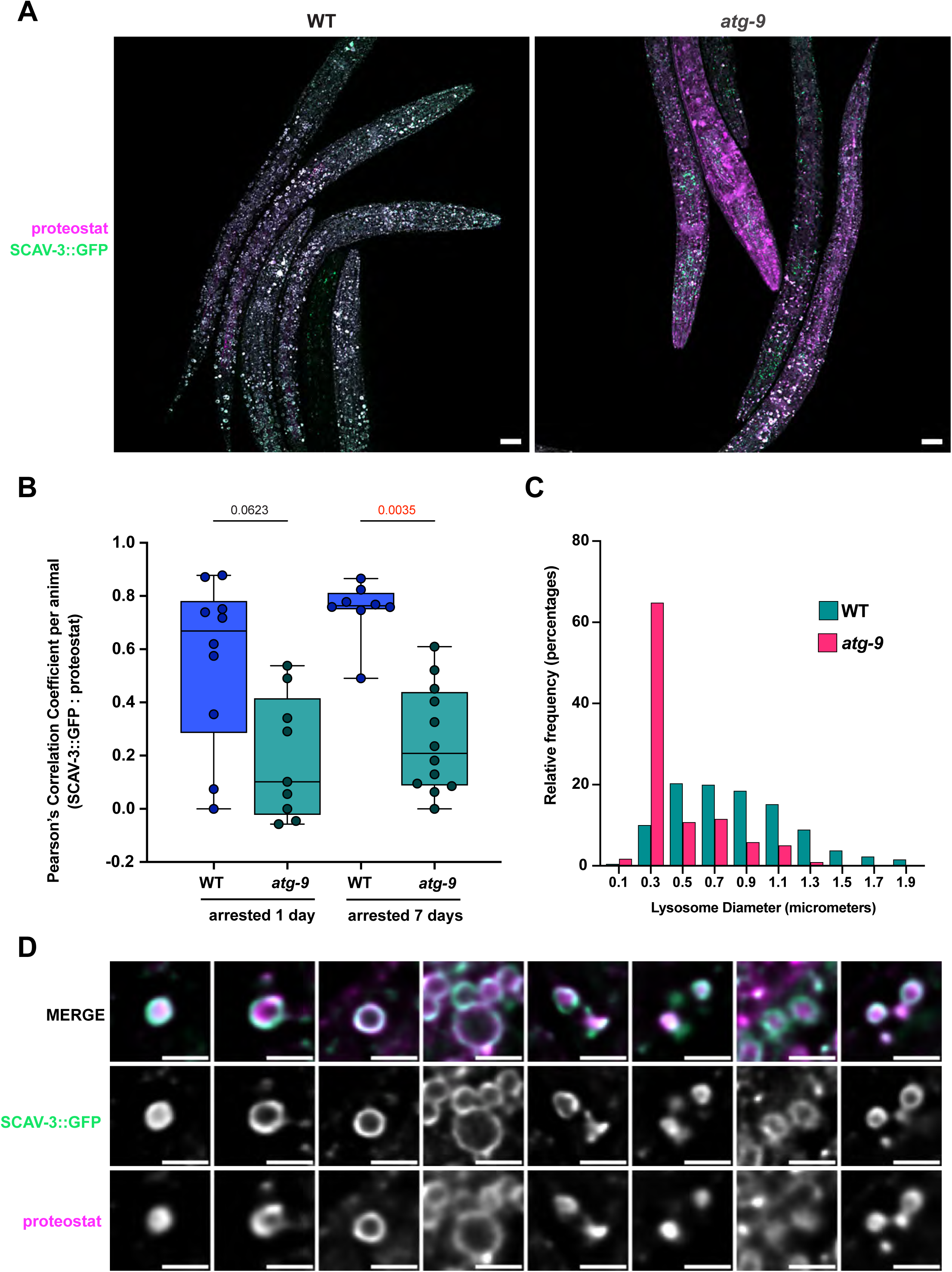
– Autophagy results in association of protein aggregates with lysosomes during L1 arrest. **(A)** Representative images of proteostat and SCAV-3::GFP marked lysosomes in WT and *atg-9* animals after five days of L1 arrest. Scale bars = 10 μm. **(B)** Pearson’s correlation coefficients of proteostat-SCAV-3::GFP signal in WT and *atg-9* animals after 1 or 7 days of L1 arrest. Pearson’s correlation coefficients were calculated for each image in a Z series encompassing an entire animal and the median Pearson’s correlation coefficient is plotted. P values were calculated using a Kruskall Wallis test. **(C)** Lysosomes are smaller in *atg-9* mutants during L1 arrest. Diameters of SCAV-3∷GFP Lysosomes were measured from animals arrested for seven days. **(D)** Representative images of SCAV-3::GFP and proteostat in WT animals L1 arrested for 5 days showing the types of associations between lysosomes and SCAV-3::GFP. Scale bars = 2 μm.

Protein aggregates that are taken into endocytic vesicles and lysosomes can cause their rupture and the recruitment of galectins, such as Galectin3, which recruits various machineries to respond to lysosomal membrane disruption ^50,51^. Galectin3 binds to galactoside sugars that are found inside of the lysosomal lumen. When lysosomes are damaged, resulting in the permeabilization of their membranes, Galectin-3 can access and bind to these sugar molecules ^48^. We created a monomeric Azami Green (mAG) fusion to human Galectin3 driven by the constitutive promoter *eft-3* and expressed from a single copy insertion at a safe harbor locus to assess whether lysosomes become damaged during L1 arrest ^52^. mAG::Galectin3 puncta were rare in un-arrested L1 animals, but were recruited to ∼ 1,500 punctate structures in various tissues of *C. elegans* early during L1 arrest, indicating lysosomal damage (**Figure 4A-B**). However, damage to lysosomes in L1 arrested animals did not lead to a complete leakage of lysosomal enzymes that may cause cell death. In hypodermal cells, NUC-1::mCherry, a lysosomal nuclease, was retained in lysosomes marked by mAG::Galectin3 (**Figure 4C**). Consistent with autophagy contributing to lysosomal damage during L1 arrest, fewer mAG-Galectin3 puncta were observed in *atg-9* mutants (**Figure 4B**). mAG::Galectin3 was localized in puncta on the edges of spherical proteostat structures, presumably lysosomes, suggesting that it is recruited to localized damage on lysosomes that may be caused by protein aggregates (**Figure S3A**).

**Figure 4.**
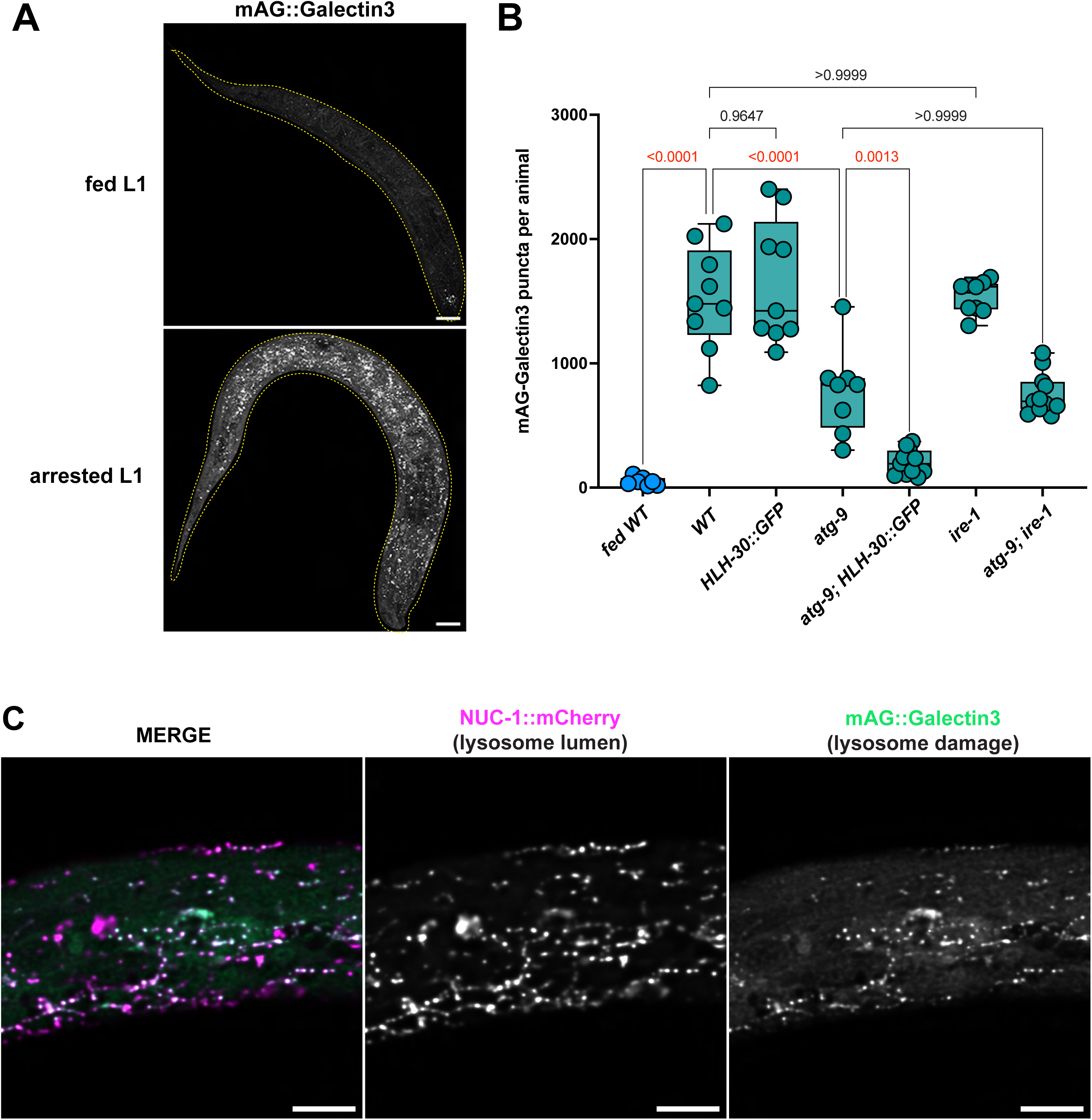
– Autophagy contributes to lysosome damage during L1 arrest. **(A)** Representative images of mAG::Galectin3 in un-arrested and L1 animals arrested for one day. Images are a maximum intensity project of the entire animal. Dashed yellow lines show the outline of the animals. Scale bars = 10 μm. **(B)** Quantification of mAG-Galectin3 puncta in animals of the indicated genotype after two days of L1 arrest. P values are indicated in red and are from Tukey’s multiple comparison test of an ordinary one-way ANOVA. **(C)** mAG::Galectin3 puncta are associated with lysosomes. L1 animals arrested for 7 days and expressing mAG::Galectin3 and NUC-1::mCherry (specifically in hypodermal cells) were imaged. A single focal plane is depicted. Scale bars = 5 μm.

As combined autophagy inhibition and HLH-30::GFP overexpression almost completely restores PTLA in *xbp-1* mutants, we tested if these combined interventions led to a further decrease in lysosome damage (**Figure 2G**). The mAG::Galectin3 reporter formed much brighter and much smaller structures than HLH-30::GFP, which mainly localized to the nucleus in L1 arrested animals, enabling us to confidently identify mAG::Galectin3 puncta: in animals expressing only HLH-30::GFP, we only identified ∼100 puncta that met the same definition as mAG::Galectin3 puncta, which formed ∼1,500 puncta in WT animals (data not shown). Consistent with HLH-30::GFP overexpression having a marginal effect on PTLA, the numbers of mAG::Galectin3 puncta were indistinguishable between WT and HLH-30::GFP overexpressing animals (**Figure 4B**). In contrast, *atg-9* mutants overexpressing HLH-30::GFP had far fewer mAG::Galectin3 puncta than WT, *HLH-30::GFP*, and *atg-9* animals (**Figure 4B**). Thus, lysosome damage, possibly resulting from protein aggregates that are delivered to lysosomes by autophagosomes is well correlated with an inability of *ire-1* and *xbp-1* animals to develop after prolonged L1 arrest (**Figures 2G and 4B**).

### Autophagy causes lysosome damage in quiescent mammalian cells *in vitro*

We asked whether lysosome damage is a particular aspect of cells in L1 arrested *C. elegans* because of nutrient deprivation or if it is a feature of other mitotically quiescent cells. We could robustly induce reversible cell cycle arrest in cell culture using 3T3 Swiss Albino and C3H/10T1/2 mouse fibroblast cells and ARPE-19 human retinal epithelial cells by contact inhibition without any changes in cell culture medium such as serum withdrawal (**Figure S4A**) ^53–55^. Levels of the mitotic antigen Ki67 were almost absent in contact-inhibited cells (herein “quiescent”) but were restored 48 hours after re-plating cells at a lower cell density (**Figure S4A**). However, over time, the ability to resume proliferation in response to growth cues in 3T3 cells was diminished, with a greater proportion of cells having low Ki67 levels 48 hours after plating, as well as slower growth rates and fewer colonies formed from equivalent numbers of plated viable cells, indicating that an increasing proportion of cells transition from quiescence to irreversible cell cycle arrest (**Figure S5A-C**).

We tested whether lysosomes harbor increased amounts of protein aggregates in quiescent mammalian cells stably expressing EGFP-Tmem192, which localizes to lysosomal membranes, and stained with proteostat dye. In all cell lines tested, a greater percentage of lysosomes harbored protein aggregates in quiescent cells than in their proliferating counterparts (**Figure 5A-D**).

**Figure 5.**
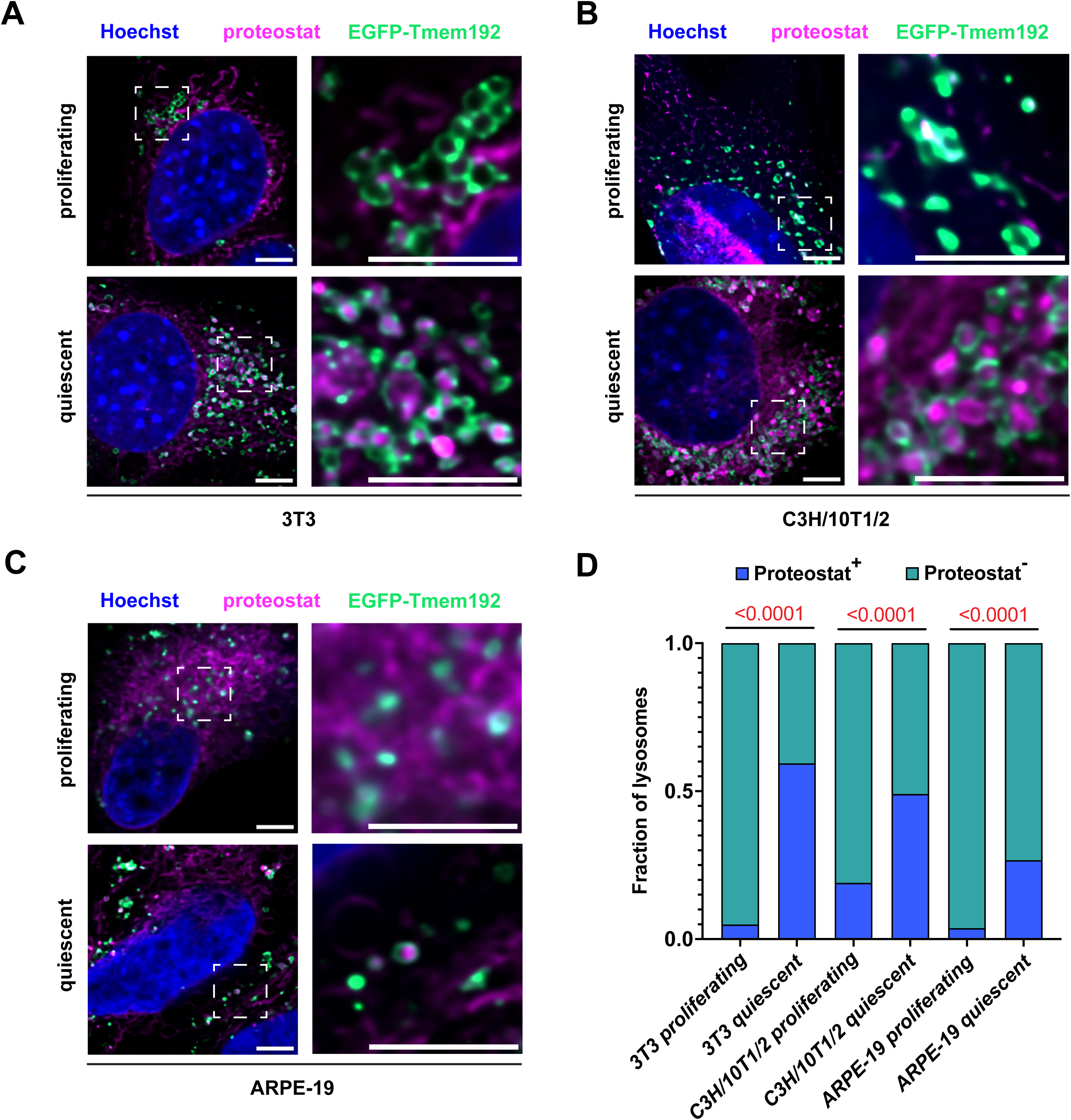
– Protein aggregates are associated with lysosomes in quiescent mammalian cells. **(A-C)** Intralysosomal protein aggregates accumulate in quiescent mammalian cells. 3T3 Swiss Albino, C3H/10T1/2 and ARPE-19 cells stably expressing EGFP-Tmem192 were grown to confluency and maintained in a contact inhibited state for 10 days. Quiescent cells, and proliferating cells as a control, were stained with proteostat to detect protein aggregates side-by-side and imaged as described in “Methods”. Depicted are representative examples of cells from each condition. Scale bars = 5 μm. **(D)** A greater percentage of lysosomes contain protein aggregates in quiescent 3T3 Swiss Albino, C3H/10T1/2 and ARPE-19 cells than their proliferating counterparts. P values were calculated using Chi-squared to test for differences in the proportion of lysosomes containing protein aggregates (proteostat+) and those that did not (proteostat-) in proliferating versus quiescent cells.

To assess whether lysosomes become damaged in quiescent mammalian cells, we again utilized the Galectin3 reporter of lysosomal damage. We found that mAG-Galectin3 is primarily localized to the cytoplasm and nucleoplasm in proliferating cells but is strikingly recruited to puncta mainly associated with lysosomes in all quiescent cell lines tested (**Figure 6A-D**). Thus, lysosome damage, possibly because of protein aggregates, is a hallmark of quiescent cells.

**Figure 6.**
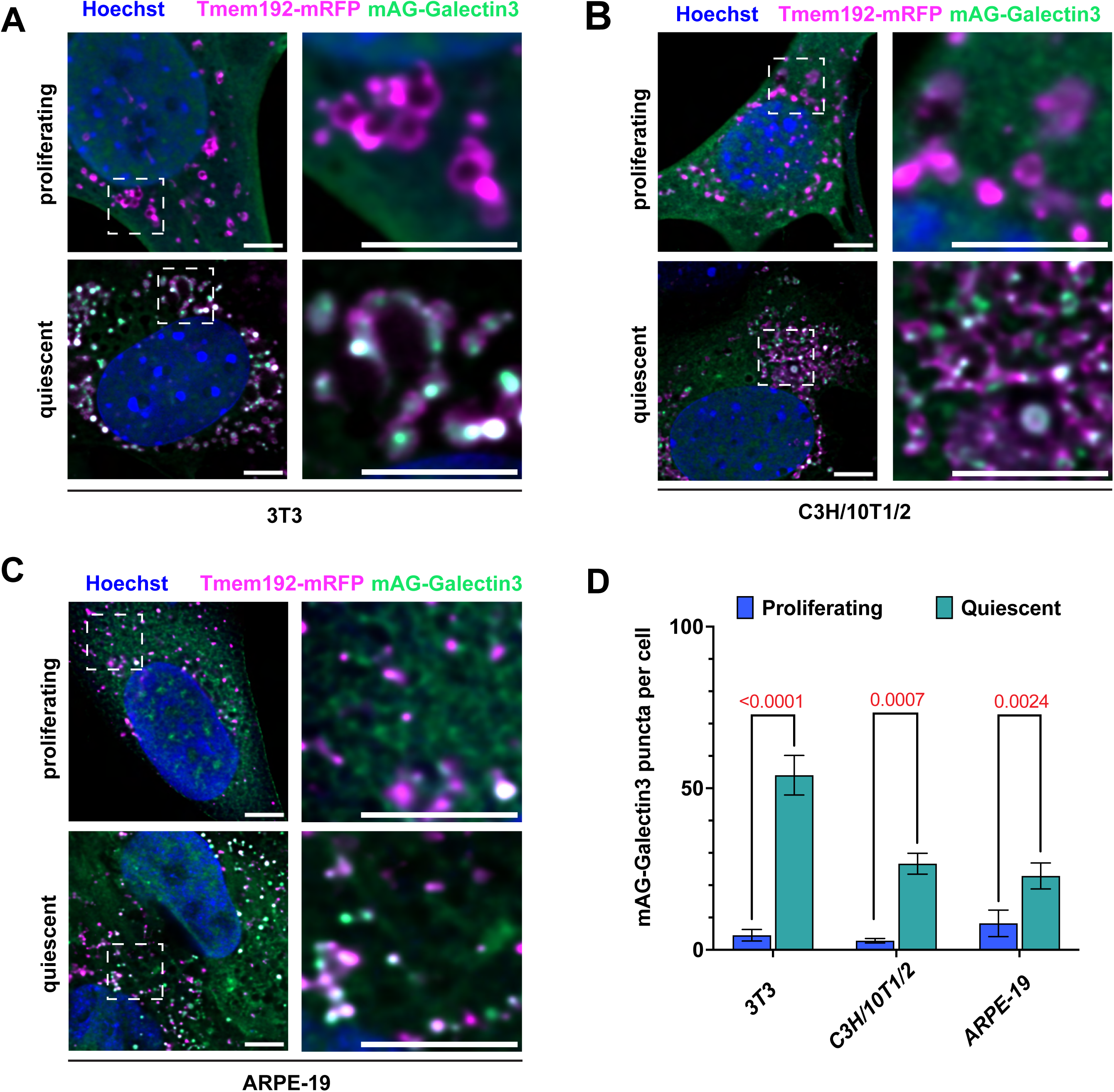
– Lysosome damage accumulates in quiescent mammalian cells. **(A-C)**. 3T3-Swiss Albino, C3H/10T1/2 and ARPE-19 cells stably expressing mAG-Galectin3 and Tmem192-mRFP1 were grown to confluency and maintained in a contact inhibited state for 10 days. Quiescent cells and their proliferating counterparts were fixed, stained with Hoechst and images. Scale bars = 5 μm **(D)** Quiescent cells contain a greater number of mAG-Galectin3 puncta than their proliferating counterparts. A single focal plane in the center of cells was used to count the number of mAG-Galectin3 puncta per cell in proliferating and quiescent cells. It was difficult to accurately segment individual contact inhibited cells, so the number of mAG-Galectin3 puncta was divided by the number of cells in the image, and this number was calculated from four fields of view. Plotted values represent mean ± SEM.

We used CRISPRi to assess the role of autophagy in lysosomal damage in quiescent 3T3 cells. Using an autophagy flux reporter, we confirmed that CRISPRi of *Atg5*, *Atg7* and *Atg9a* inhibited autophagy in response to mTOR inhibition (**Figure S6A**). Knockdown of autophagy genes reduced the association of mAG-Galectin3 with lysosomes and the proportion of lysosomes containing protein aggregates (**Figure 7A and B**). Thus, consistent with our findings in *C. elegans*, autophagy contributes to lysosome damage in quiescent mammalian cells possibly by contributing to the accumulation of protein aggregates within lysosomes.

**Figure 7.**
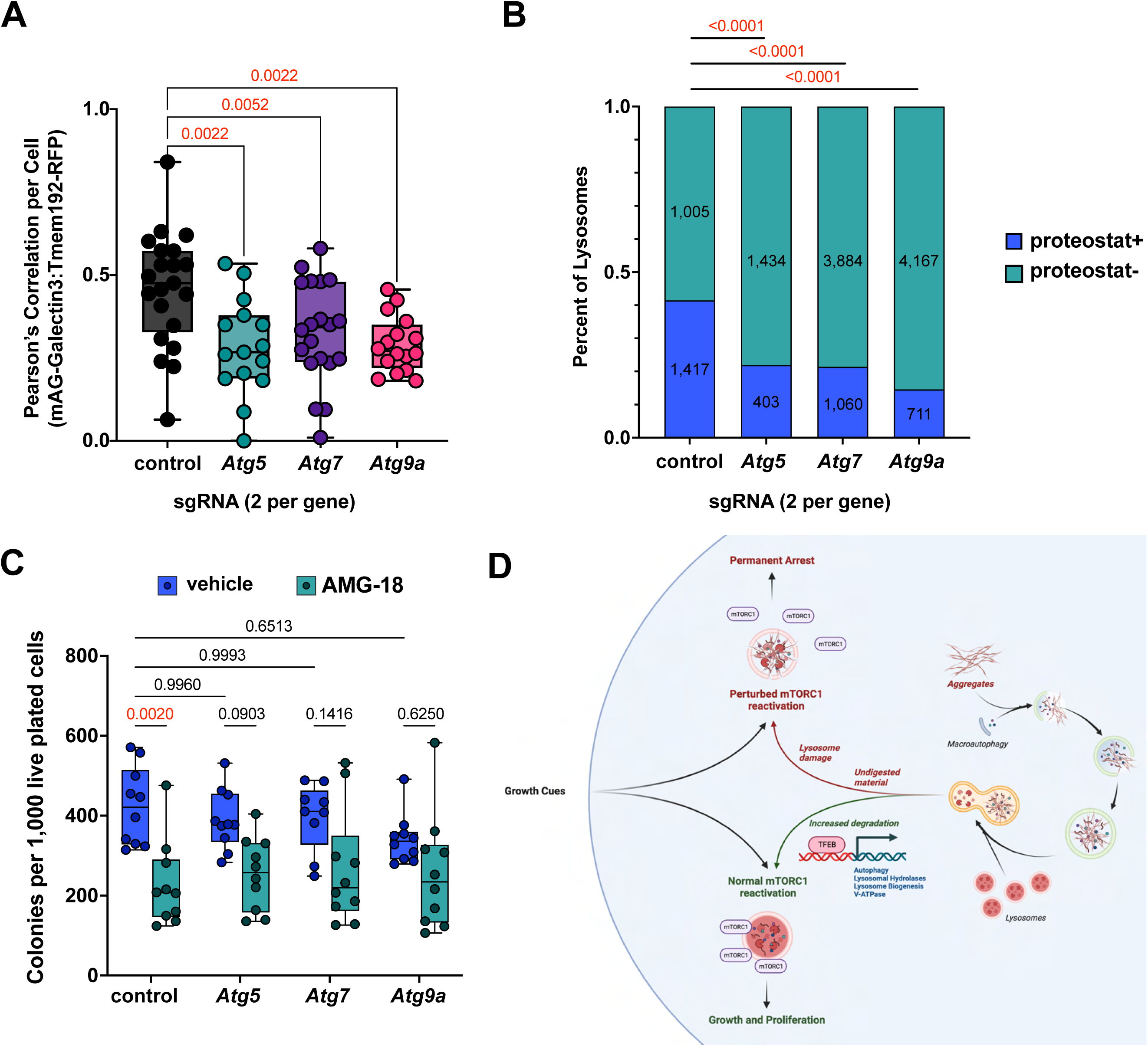
– Autophagy inhibition reduces lysosomal protein aggregates and lysosome damage in quiescent mammalian cells. **(A)** Knockdown of autophagy genes reduces localization of mAG::Galectin3 with Tmem192::mRFP lysosomes. CRISPRi was used to knock-down Atg5, Atg7 and Atg9a in cells expressing mAG::Galectin3 and Tmem192-mRFP. Cells were plated at a high density and kept in a contact-inhibited state for 10 days. Cells were fixed and imaged, and the median Pearson’s correlation coefficient from Z-series of cells expressing are plotted. Adjusted P-values were calculated from Tukey’s multiple comparisons test of an ordinary one-way ANOVA. **(B)** Knockdown of autophagy genes reduces the percentage of lysosomes containing protein aggregates. CRISPRi was used to knock-down Atg5, Atg7 and Atg9a in cells expressing EGFP-Tmem192. Cells were plated at a high density and kept in a contact-inhibited state for 10 days. Cells were fixed and stained for proteostat as described in “Methods” then imaged. P values were calculated using Chi-squared to test for differences in the proportion of lysosomes containing protein aggregates (proteostat+) and those that did not (proteostat-) in control cells and autophagy knockdown cells. In-laid numbers represent the total number of lysosomes in each group. **(C)** Knockdown of autophagy genes mitigates defective colony formation from AMG-18 treated 3T3 cells. Cells were grown to confluency, then 5 μM AMG-18 (or DMSO as a vehicle control) was added and replenished every other day for 10 days. Cells were trypsinized and 1,000 live cells were plated and allowed to form colonies for seven days. Each point represents an independent biological replicate pooled from two guide RNAs per gene. P values were calculated using Tukey’s multiple comparisons test from a two-way ANOVA. **(D)** Model of lysosome damage in quiescent cells.

We also used CRISPRi and a chemical inhibitor of Ire1α to assess the role of autophagy and the UPR in re-activation of quiescent mammalian cells. We grew 3T3 cells to confluency, then inhibited Ire1α using the potent and specific inhibitor AMG-18 (Kira8) for 10 days ^56,57^. We subsequently split cells and plated 1,000 live cells on plates and assessed the number of colonies formed one week later. We found that knockdown of autophagy genes did not significantly affect the number of colonies formed from quiescent cells replated after 10 days of arrest (**Figure 7C**). Treatment of quiescent cells with AMG-18 significantly reduced subsequent colony formation, and inhibiting autophagy mitigated the decrease in colonies that are formed from cells treated with AMG-18: treatment with AMG-18 reduced colony formation by 46% in control cells, but by only 35%, 33% and 25% in *Atg5*, *Atg7*, and *Atg9* knockdown cells, respectively (**Figure 7C**). Thus, consistent with our findings in *C. elegans*, reducing lysosome damage caused by autophagy in quiescent mammalian cells might be a means of boosting their re-activation potential during aging when function of the UPR declines.

## Discussion

We have identified a set of processes that contribute to the declining plasticity of quiescent cells, centered around lysosomal dysfunction (**Figure 7D**). Quiescent cells have lower anabolic demands and activate autophagy during arrest partly through reduced mTORC1 activity. This contributes to an increase in lysosome damage, likely through an excessive delivery of protein aggregates that are known to damage lysosomal membranes. This process can be slowed or reduced in a synergistic manner by inhibiting autophagy during arrest and by boosting lysosome biogenesis or function. A degree of cell growth being required for cellular proliferation, re-activation of quiescent cells requires mTORC1 activation, but this is likely inefficient in cells with damaged lysosomes.

Autophagy is generally thought to be a process that promotes longevity by the turnover of dysfunctional and damaging constituents of cells. Consistent with this view are the observations that autophagy is required for the success of all known longevity paradigms in *C. elegans*, an organism that does not possess adult somatic stem cells, and that adult-onset perturbation of autophagy causes premature death in mice ^58^. Likewise, flux through the autophagy pathway declines with age, and longevity paradigms tend to mitigate this decline ^5,7,59^. Together these findings suggest that enhanced autophagy may promote longevity. However, our findings suggest that autophagy may also have a detrimental effect on quiescent cells that make important contributions to tissue homeostasis and repair. Likely through the delivery of protein aggregates to lysosomes, autophagy contributes to increasing lysosomal damage during quiescence, leading to impaired mTORC1 re-activation and likely other effects on cellular homeostasis. Our data indicate that lysosome damage is a feature of quiescent cells that transition towards irreversible growth arrest, both in *C. elegans* and mammalian cells. Our study adds to previously published findings centering lysosomal function between reversible cell cycle arrest (quiescence) and a transition towards irreversible cell cycle arrest (senescence) ^5,60–62^. It will be important to examine in future studies how lysosome damage is regulated in quiescent cells *in vivo*.

The accumulation of lysosomal damage in quiescent cells is somewhat perplexing given the numerous means cells utilize to repair, degrade, and replace damaged lysosomes ^48,63,64^. Most mechanistic studies of lysosomal damage and repair processes have used acute treatments to cause damage to lysosomes in proliferating cells. It may be that proliferative growth cues are required for the repair of damaged lysosomes or that the dilution of protein aggregates or damaged lysosomes through cell growth and division is an important factor influencing the accumulation of lysosome damage. Lysosome damage in quiescent cells constitutes a new paradigm to explore lysosome repair processes and lysosomal stress responses, warranting future study.

What is the connection between IRE-1 and XBP-1 and lysosome damage? Although interventions that reduce lysosome damage correlate well with improved recovery from quiescence, as shown in Figure 4B, *ire-1* mutants did not have increased lysosomal damage compared to WT animals and lysosomal damage occurs prior to when L1-arrested *C. elegans* and contact-inhibited quiescent cells completely lose the ability to develop or resume proliferation. This suggests that lysosomal damage *per se* does not contribute to permanent cell cycle arrest. Instead, we hypothesize that IRE-1 and XBP-1 play a role in the response to lysosomal damage that is required to prevent permanent cell cycle arrest. One possibility is that re-generation and/or repair of lysosomes requires an optimally functioning ER, since lysosomal proteins are first synthesized in the ER. However, given the numerous genes and processes regulated by IRE-1 and XBP-1, further study is needed to systematically identify the processes performed by IRE-1 and XBP-1 that respond to lysosome damage and contribute to mTORC1 re-activation and cell cycle re-entry.

Our findings differ in some ways from those of earlier studies studying quiescent adult stem cells *in vivo*. The first found that that *Atg7^-/-^*muscle stem cells have increased rates of cell senescence and was interpreted to indicate that autophagy prevents senescence by removing dysfunctional mitochondria ^6^. A recent study has clarified the role of autophagy in hematopoietic stem cells in mice, finding that autophagy is essential for the maintenance of hematopoietic stem cells and their multipotent progenitors ^67^. Knockout of *Atg5* or *Atg7* in hematopoietic stem cells led to increased amino acid uptake and hyperactivation of mTORC1, and defects in hematopoietic stem cell functions were partially reversed by rapamycin treatment ^67^. However, the findings from these studies often rely on cell growth from transplantation experiments and cannot exclude the role of reduced growth rates of autophagy deficient cells. It may be that a decrease in cell proliferation rates in *Atg5^-/-^* and *Atg7^-/-^* cells might appear as increased proportion of senescent stem cells and perturbed outgrowth of transplanted cells ^65^. Our study tested for an absolute ability to exit quiescence, either in the ability of *C. elegans* to develop following L1 arrest or in the ability of cells to form colonies. These studies did not explore the role of the UPR, and our findings indicate that the deleterious effects of autophagy-mediated lysosome damage become more apparent in UPR-deficient cells. Although the discrepancies between our results and these studies might simply be explained by different requirements for autophagy-mediated process in certain cell types or experimental paradigms, alternative explanations might also exist. For example, the Atg8/LC3 lipidation machinery, which includes Atg5 and Atg7, is required for Tfeb activation and nuclear translocation following acute lysosomal damage ^66^. Although *Atg5* and *Atg7* knockout cells would have less lysosome damage resulting from autophagy, other processes also contribute to lysosome damage in cells, such as the cellular uptake of proteinaceous aggregates. The *Atg5* and *Atg7* knockout cells utilized in the above studies therefore may be deficient in activation of Tfeb that responds to lysosome damage that we have identified as a conserved feature of many types of quiescent cells. Therefore, defects in these cells may stem from perturbed responses to lysosomal damage, not from defects in autophagy *per se*. Considering our findings, the role of autophagy and lysosome damage on numerous types of quiescent cells *in vivo* should continue to be examined.

Although our findings indicate that autophagy causes lysosome damage in quiescent cells, given the numerous roles autophagy plays in cellular and organismal homeostasis and in the maintenance of certain stem cell pools, inhibiting autophagy at the organismal level is certain to be detrimental overall and therefore unlikely to be a good therapeutic strategy to protect quiescent cells in mammals ^67^. Indeed, whole body of knockdown of *Atg7* causes death about 30 days after treatment in mice ^68^. Furthermore, as discussed below, other processes beyond autophagy can contribute to lysosome damage, so lysosome damage would not be completely eliminated by reduced autophagy. Our findings point instead to other interventions that might maintain or restore the appropriate re-activation of quiescent cells in age. First, interventions that specifically promote lysosomal function and reduce lysosomal damage in quiescent cells should boost their regenerative potential. As suggested by our findings, common ways of boosting lysosomal function, such as activation of TFEB, have a negative consequence of boosting autophagy through the transcriptional upregulation of core autophagosome machinery. Transcriptional programs or other paradigms that specifically boost lysosome function or make them more resistant to the effects of damaging agents such as protein aggregates may be a better intervention for quiescent cells ^69^. Second, the IRE-1/XBP-1 branch of the UPR seems to have a critical role in responding to lysosomal damage and allowing for a robust re-activation of mTORC1 and anabolic cellular growth. Since this pathway becomes dysfunctional with age in some cells and tissues, finding ways to mitigate or bypass this decline may prove useful ^70–73^. Finally, direct activation of mTORC1 might be a means of bypassing the perturbed lysosomal homeostasis of quiescent cells to facilitate cell growth and proliferation ^13^. Indeed, the small molecule MHY1485, a putative activator of mTORC1 and autophagy inhibitor, was found to improve the regeneration of retinal pigment epithelium following injury in Zebrafish ^74^. Thus, an acute, pharmacological activation of mTORC1 specifically following injury may stimulate a restoration of tissue homeostasis, yet direct, specific activators of mTORC1 are not widely utilized or explored.

Lysosomal accumulation of protein aggregates, and resulting lysosomal damage, is likely to play a role in cellular dysfunction in other types of quiescent cells and in human diseases. Previous studies have found that protein aggregates are associated with lysosomes in quiescent neural stem cells and senescent cells ^5,62,75^. Although the targeting of protein aggregates to lysosomes by autophagy can cause lysosome damage, other processes are likely also involved. Lysosome disruption caused by endocytosis or macropinocytosis of human disease associated proteinaceous aggregates/fibrils, such as Aβ_1-42_, Tau and α-synuclein may also be a means for protein aggregates to spread within tissues ^50,76–80^. Thus, finding ways to mitigate endolysosomal damage from protein aggregates may prove useful for healthy aging of various cell types and tissues in humans, including in neurodegenerative diseases, where it may both prevent the spread of protein aggregates within the brain and stimulate the regenerative potential of quiescent neural stem cells ^5,77^. The striking accumulation of damaged endolysosomes in quiescent cells that we have uncovered may be a useful paradigm for future exploration of interventions that increase the resiliency of the endolysosomal system in quiescent cells and beyond, paving the way towards treatments for age-associated diseases associated with endolysosomal dysfunction.

## Acknowledgements

We thank the University of California, Berkeley cell culture facility for providing cell lines and the Vincent J. Coates Genomics Sequencing Lab for sequencing of *C. elegans* genomic sequences. AM was supported by a fellowship from the Damon Runyon Cancer Research Foundation; KW was supported by a fellowship from the National Science Foundation; AD is an investigator of the Howard Hughes Medical Institute.

## Author Contributions

Conceptualization, AM and AD; Methodology, AM; Validation, AM; Formal Analysis, AM; Investigation AM, AP, SL, AL, KW, EK, JD, LJ; Resources Provision, AD; Data Curation, AM, LJ and AD; Writing – Original Draft Preparation, AM; Writing – Review & Editing, AM, AP, SL, AL, KW, EK, JD, LJ, AD; Visualization Preparation, AM. Supervision, AD; Project Administration, AM; Funding Acquisition AM and AD.

## Supplemental Information

Document S1. Figures S1-6

## Supplemental Figure Legends

**Figure S1 – *ire-1* animals can ingest food after prolonged L1 arrest.** (A) WT and *ire-1* animals were subject to a seven-day L1 arrest, then fed *E. coli* OP50 expressing RFP for eight hours. Animals were paralyzed with 100 mM NaN_3_ and then imaged. In control experiments, animals were plated on OP50 and scored for their ability to develop to the at least the L3 larval stage after 2-4 days. Scale bars = 10 um.

**Figure S2 – SQST-1, but not PDR-1 or EPG-7, influences PTLA.** (A-C). Animals of the indicated genotypes were subjected to a five-day L1 arrest, then assessed for viability and their ability to develop to at least the L3 larval stage after 2-4 days with food. Each symbol represents an independent biological replicate. P values were calculated from an ordinary Two-Way ANOVA with Tukey’s multiple comparisons test.

**Figure S3 – mAG::Galectin3 puncta are often associated with proteostat in L1-arrested animals.** (A) Animals expressing mAG::Galectin3 were subjected to L1 arrest for three days, then stained for proteostat and imaged as described in Methods. Scale bars = 2 um.

**Figure S4 – Reversible cell cycle exit in 3T3 Swiss Albino, C3H10T1/2 and ARPE-19 cells**. Proliferating cells are cells that were maintained in sub confluent (<70% confluency) for two weeks after thawing before harvesting and staining for Ki67 labeling. Quiescent cells were harvested for Ki67 labeling 4 days after reaching confluency. Re-activated cells are quiescent cells that were passaged 1:10 (3T3 and C3H10T1/2) or 1:5 (ARPE-19) and harvested for Ki67 labeling 48 hours later. Unlabeled cells are pooled proliferating, quiescent and re-activated cells that were not stained for Ki67. Cells were labeled with Ki67 antibodies and analyzed by flow cytometry as described in Methods.

**Figure S5 – Quiescent 3T3 Swiss Albino cells have diminished re-activation over time** (A) 3T3 Swiss Albino cells were grown to confluency and maintained for the indicated number of days. Cells were split 1:10 and harvested 48 hours later, stained for Ki67 and analyzed by flow cytometry. (B) 3T3 Swiss Albino cells were grown to confluency and maintained for the indicated number of days. They were then split and 5,000 cells per cm^2^ were plated six well plates and imaged on an IncuCyte and analyzed for confluency every four hours. (C) 3T3 Swiss Albino cells were grown to confluency and maintained for 45 days. Cells were passaged alongside a sub confluent culture and 1,000 cells were plated on 10 cm dishes and incubated for seven days until cell colonies formed. Cells were fixed with 4% PFA and stained with Crystal Violet.

**Figure S6 – CRISPRi against *Atg5, Atg7, and Atg9a* reduces autophagy flux.** (A) Cells expressing mCherry-EGFP-LC3B, dCas9 and one of two sgRNA constructs against the indicated genes, or targeting no genes as a control, were treated with 100 nM Torin1 or vehicle (DMSO) and imaged every 30 minutes on an IncuCyte live-cell analysis instrument. Vehicle control cells were “no target” control cells.

## STAR Methods

### Key Resources Table

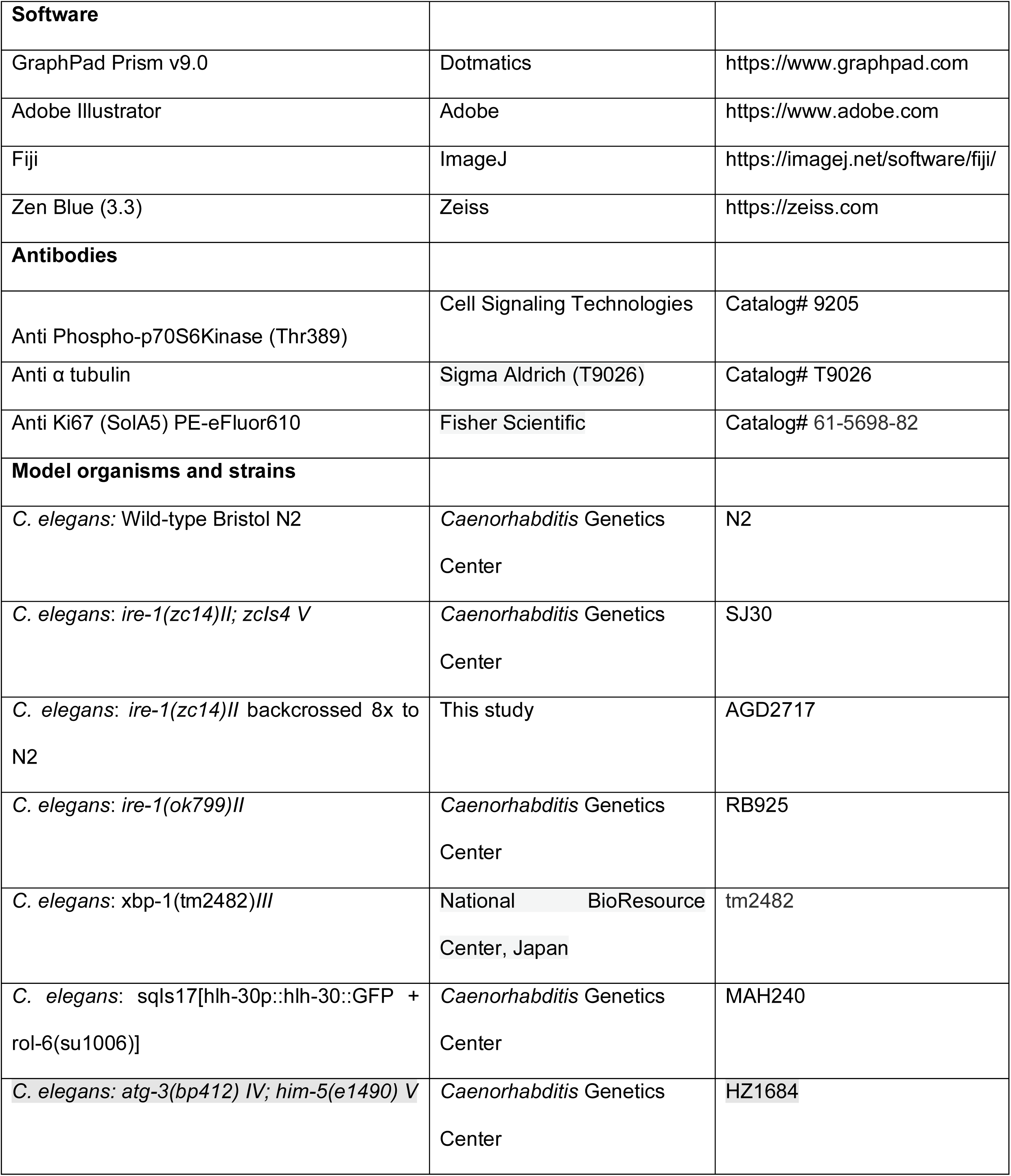
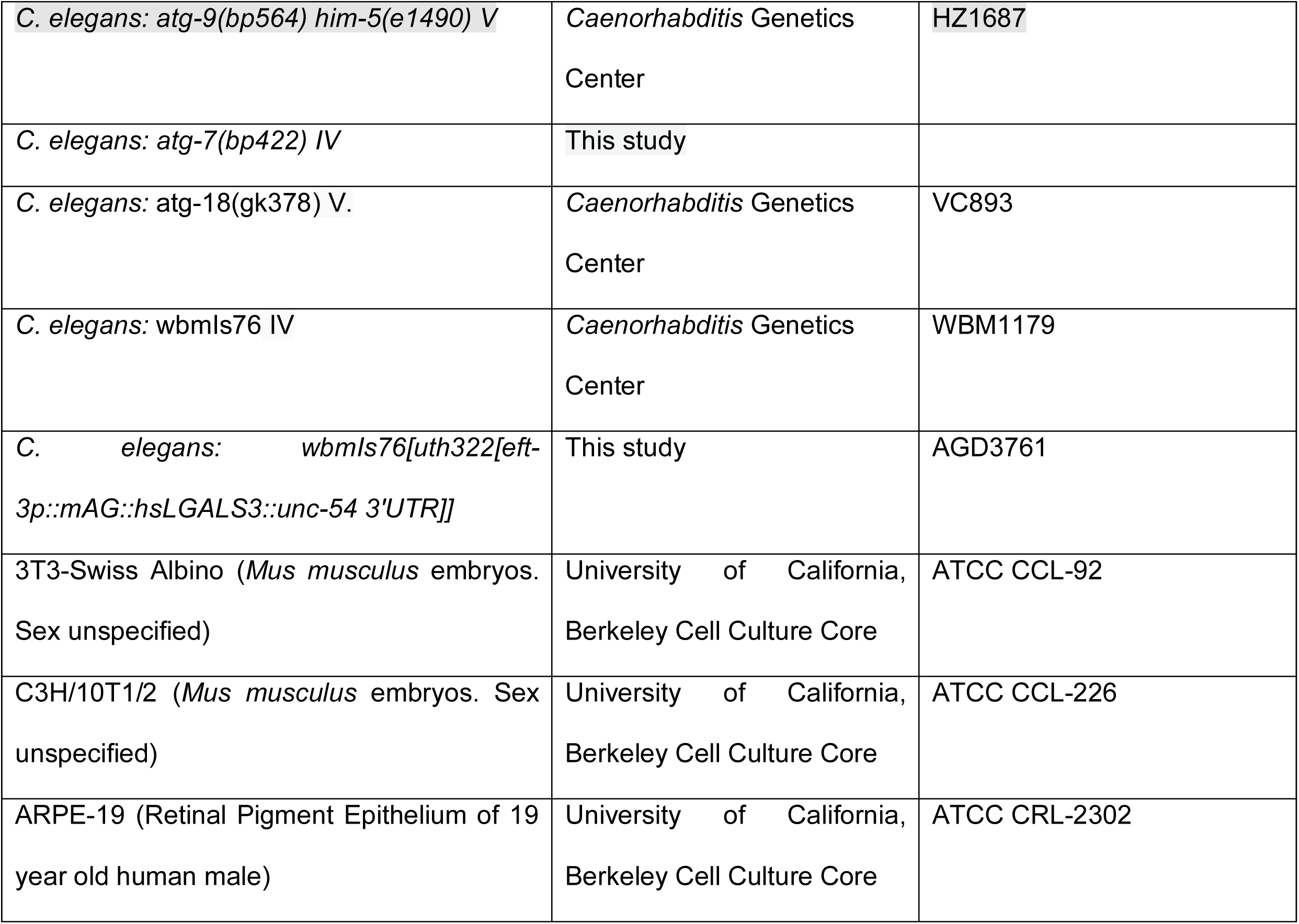

### Resource availability

#### Lead contact

Further information and requests for resources and reagents should be directed to and will be fulfilled by the lead contact, Andrew Dillin

#### Materials availability

This study generated unique reagents, see “Key Resources Table”

#### Data and Code Availability

No original code was used in this study.

### Experimental Models

#### C. elegans

*C. elegans* WT Bristol N2 were used as the WT strain background. All mutant strains were backcrossed to N2 at least six times before analysis. Animals were maintained on NGM agar plates seeded with *E. coli* OP50 bacteria.

#### Mammalian Cells

3T3-Swiss Albino (ATCC CCL-92), C3H/10T1/2 (ATCC CCL-226) and ARPE-19 (ATCC CRL-2302) cells were obtained from the University of California, Berkeley cell culture core facility and were free of mycoplasma contamination. Cultures remained free of mycoplasma contamination as assessed by periodic Hoechst staining.

All cells were grown in Human Plasma-Like Medium (HPLM; Gibco) + 10% Fetal Bovine Serum (FBS) + penicillin/streptomycin in a humidified 37 C incubator in 5% CO_2_ and atmospheric O_2_ in cell-culture treated plastic dishes (Corning) for routine culturing and growth-based assays or on glass-like polymer dishes (CellVis) for imaging. Media was replaced every other day for proliferating and quiescent cells ^86^.

### Method Details

#### L1 arrest

Eggs were harvested from well-fed adult hermaphrodites by bleaching. Briefly, animals were washed off plates with M9 buffer, and collected by centrifugation at 1,400 g for 30 seconds in a 15 mL conical tube. Animals were treated with bleaching solution (1.5 % sodium hypochlorite + 645 mM KOH) for about 5 minutes until the adults had broken apart. The eggs were harvested by centrifugation and washed 5x with 15 mL of M9 buffer. Eggs were vortexed for about 1 second and then resuspended in S-basal medium (5.85 g/L NaCl, 1 g/L K_2_ HPO_4_, 6 g/L KH_2_PO_4_, 5 mg/L cholesterol (from a 5 mg/ml stock in ethanol)) at 5-7 eggs per ul. One day of L1 arrest was defined as 24 hours after bleaching.

To assess animal viability and development, 150-250 animals were spotted directly onto the lawn of OP50 bacteria of 60 mm NGM plates. After 1 hour, animals that had not moved were deemed dead. After 2 days, animals developing to the L3-L4 stage were picked off plates and counted. This was repeated 3 and 4 days after plating and number of animals removed from plates on days 2-4 was totaled.

#### EMS mutagenesis screen

Approximately 300 *ire-1(zc14)* or *ire-1(ok799)* L4 larvae were picked into M9 and washed once with M9. Animals were resuspended in M9 buffer with 0.5% of EMS (methanesulfonic acid, ethyl ester, Sigma #M-0880). The tube was nutated at 20 C for 4 hours. After incubation, P0 animals were washed four times with 1mL M9 and allowed to recover on plates. The next day, mutagenized worms that survived EMS treatment were picked onto 10 cm plates with OP50 (10 per plate) and allowed to lay eggs for 48 hours. Three days later, gravid adult F1 animals were bleached and resulting F2 animals were subjected to L1 arrest for 14 days at 20 C. F2 animals were then plated on large OP50 plates, and after 2-4 days at 20 C animals that had developed to the L4 stage were picked singly to OP50 plates at 20 C. Many F2 animals were sterile and were unable to establish lines. Established lines were re-tested for their ability to develop after prolonged L1 arrest.

Purification of genomic DNA from established suppressor lines was performed using the Puregene Cell and Tissue Kit (Qiagen), as previously described ^82,83^. 2ug of purified DNA was sheared using a Covaris S220 focused-ultrasonicator to produce ∼400 bp fragments. Library preparation was performed with 1ug of sheared DNA using Kapa Biosystems Hyper Prep Kit (Roche, product number KK8504) dual index adapters (KAPA, product number KK8727). Sequencing was performed using the Illumina NovaSeq6000 platform through the Vincent J. Coates Genomic Sequencing Core at University of California, Berkeley. Raw reads were uploaded to the Galaxy project web platform and the public server was used to analyze the data ^84^. Reads were aligned using the Bowtie2 tool with WBcel235/ce11 as the reference genome. The MiModD tool suite was used on the Variant Allele Contrast (VAC) mapping mode to call, extract and filter variants to compare mutants to the parental, un-mutagenized strain ^85^.

#### Creation of mAG∷Galectin3 transgenic *C. elegans*

Animals expressing the mAG∷Galectin3 transgene were created using the SKI-LODGE method ^81^. Briefly, mAG-Galectin3 was PCR amplified from Addgene plasmid #62734 and purified by ethanol precipitation. The repair template, as well as purified Cas9 ribonucleoprotein complexes with *dpy-10* crRNA were injected into the gonads of D1 adult WBM1179 animals using the SKI-LODGE system as previously described ^81^. Lines of fluorescing animals were then genotyped by sequencing analysis to ensure correct insertion of the mAG∷Galectin3 transgene.

#### Lentivirus production and stable cell line creation

The following guide RNAs were cloned in pLENTIGUIDE F-E, which was modified from lentiguide-PURO with the F-E modification for increased potency used for CRISPRi ^87,88^. pHR-UCOE-SFFV-Zim3-dCas9-P2A-Hygro was a gift from Marco Jost & Jonathan Weissman (Addgene plasmid # 188768; http://n2t.net/addgene:188768; RRID:Addgene_188768), mAG-GAL3 was a gift from Niels Geijsen (Addgene plasmid # 62734; http://n2t.net/addgene:62734; RRID:Addgene_62734) ^52,89^.

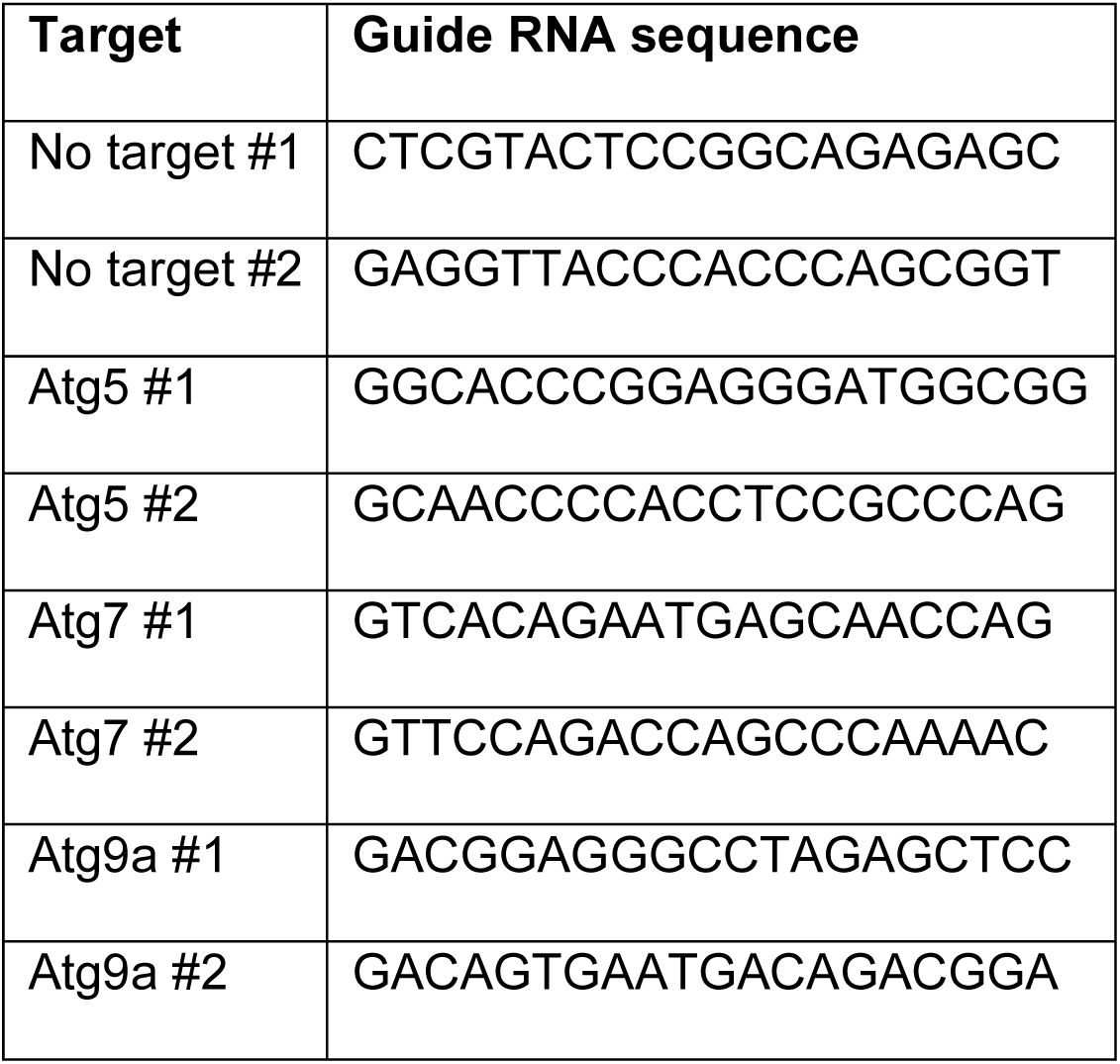

#### Western Blots

100,000 L1 animals were harvested by centrifugation at 1,400 g for 30 seconds in a 15 mL conical tube and washed twice with M9 buffer. Animals were centrifuged once more without additional M9 to collect residual buffer in the bottom of the tube, which was then aspirated. 100 ul of Urea Lysis Buffer (8 M Urea, with 1x NuPage LDS sample buffer + 1% PMSF + 1% phosphatase inhibitor cocktail) was added to animals, which were then transferred to a 1.5 mL microfuge tube and flash frozen in liquid nitrogen and stored at −80C. To harvest un-arrested L1 animals, adult animals and larvae were thoroughly washed off of OP50 plates, leaving behind eggs that hatched as L1 larvae harvested by centrifugation as described above 12 hours after adults and larvae were removed from plates. Arrested animals that were subsequently fed were concentrated by centrifugation and spotted onto OP50 plates and harvested by centrifugation 12 or 24 hours later as described above.

Samples were thawed and sonicated with a probe sonicator (Qsonica Q700) with an amplitude of 10 for seven cycles of two seconds on and one second off. Beta-mercaptoethanol was added to a final concentration of 5%. Samples were then heated at 70 C for 10 minutes. Before SDS-PAGE, samples were centrifuged at 17,000 g for 2 minutes.

Samples were separated on 1.5 mm 4-12% NuPage Bis-Tris gels using 100 V for 2.5 hours, then transferred to 0.45 um nitrocellulose membranes (BioRad Laboratories) for 80 minutes at 100 V. Membranes were blocked in LiCor Intercept TBS blocking buffer for 1 hour at RT, then probed overnight with 1:1,000 pT389 S6K or 1:20,000 alpha tubulin antibody at 4 C in LiCor Intercept TBS blocking buffer. Membranes were washed three times for 5 minutes with TBS + 0.05% Tween-20, then probed with appropriate fluorescent secondary antibodies at 1:10,000 for 1 hour in TBS + 0.05% Tween-20, then washed three times for 5 minutes with TBS + 0.05% Tween-20. Membranes were imaged using LiCor Odyssey and band quantification was performed using LiCor Image Studio.

#### Fluorescence microscopy

*C. elegans* L1 larvae were harvested by centrifugation at 1,500 g for 30 seconds and fixed in an equal volume of 4% paraformaldehyde in PBS for 15 minutes at 20-25 C with gentle rotation. Animals were then washed 3x in PBS, sealed under #1.5 coverslips and imaged.

A modified protocol was used to fix/permeabilize worms to stain with proteostat. First, 25,000-50,000 animals were harvested by centrifugation at 1,500 g for 30 seconds in Eppendorf Protein lo-bind tubes. After aspiration of the supernatant, the animals were resuspended in 1 mL of 30% acetone in miliQ water and rotated gently at 20-25 C for 15 minutes in the dark. Animals were washed three times with PBS, then stained with 1:2,000 proteostat in PBS for 2 hours at RT in the dark. Animals were washed three more times with PBS, sealed under #1.5 coverslips and imaged.

Mammalian cells were grown on glass-like polymer dishes (CellVis) to the desired stage, then washed once with PBS, fixed with 4% paraformaldehyde for 15 minutes at RT, then washed 3x with PBS. To stain cells for proteostat, cells were subsequently permeabilized (PBS, 0.2% Tween-20, 2% FBS) for 15 minutes, then washed three times again. Cells were stained with 1:2,000 proteostat in PBS for 2 hours in the dark at RT. Cells were then washed once with gentle rocking for 30 minutes, stained with Hoechst and imaged.

Cells and animals were imaged on an LSM900 Airyscan2 microscope with a 63x 1.4 NA oil immersion objective. Proteostat and mAG/GFP/EGFP tagged proteins were both excited with the 488 nm laser but to negate bleed-through fluorophores emitting wavelengths between 490-515 nm and 600-700 nm were used to identify mAG/GFP/EGFP and proteostat, respectively. Three dimensional images were subjected to Airyscan Filtering strength 1.0 and subsequently deconvolved using Zeiss’ “Fast Iterative” deconvolution algorithm as part of its Zen 3.1 software.

#### Image Analysis

Co-localization measurements were performed using Zen 3.1 software. After thresholding to exclude background signal in the two channels, regions of interest were drawn to encompass cells or animals. Pearson’s correlation coefficients were determined from these regions, and the median Pearson’s correlation coefficient in a three-dimensional stack of images encompassing a cell or animal was determined.

Quantification of mAG∷Galectin3 puncta in *C. elegans* were performed using the “3D Objects Counter” plugin for ImageJ.

#### Flow Cytometry

Cells were grown to the desired stage of confluency and trypsinized (TrypLE express) for 15 minutes, then quenched with an equal volume of complete growth medium and harvested by centrifugation at 300 g for 4 minutes. Cell pellets were triturated 20 times with a P200 pipette and then fixed/permeabilized with −20 C 90% methanol for 15 minutes. Cells were washed 3x with PBS, then blocked for 30 minutes with 5% normal goat serum for 30 minutes. Fluorescently labeled Ki67 antibody was then added at 1:500 and the cells were probed for 2 hours at RT. Cells were washed 3x with Tris-Buffered Saline with 0.05% Tween-20, then analyzed on an Attune NxT Acoustic Focusing Cytometer.

### Quantification and Statistical Analysis

Statistical analyses were performed in GraphPad Prism using tests described in the Figure Legends.

## References

1. López-Otín, C., Blasco, M.A., Partridge, L., Serrano, M., and Kroemer, G. (2023). Hallmarks of aging: An expanding universe. Cell 186, 243–278. 10.1016/j.cell.2022.11.001.

2. Marescal, O., and Cheeseman, I.M. (2020). Cellular Mechanisms and Regulation of Quiescence. Dev. Cell 55, 259–271. 10.1016/j.devcel.2020.09.029.

3. van Velthoven, C.T.J., and Rando, T.A. (2019). Stem Cell Quiescence: Dynamism, Restraint, and Cellular Idling. Cell Stem Cell 24, 213–225. 10.1016/j.stem.2019.01.001.

4. Kalamakis, G., Brüne, D., Ravichandran, S., Bolz, J., Fan, W., Ziebell, F., Stiehl, T., Catalá-Martinez, F., Kupke, J., Zhao, S., et al. (2019). Quiescence Modulates Stem Cell Maintenance and Regenerative Capacity in the Aging Brain. Cell 176, 1407–1419.e14. 10.1016/j.cell.2019.01.040.

5. Leeman, D.S., Hebestreit, K., Ruetz, T., Webb, A.E., McKay, A., Pollina, E.A., Dulken, B.W., Zhao, X., Yeo, R.W., Ho, T.T., et al. (2018). Lysosome activation clears aggregates and enhances quiescent neural stem cell activation during aging. Science (80-.). 359, 1277–1283. 10.1126/science.aag3048.

6. García-Prat, L., Martínez-Vicente, M., Perdiguero, E., Ortet, L., Rodríguez-Ubreva, J., Rebollo, E., Ruiz-Bonilla, V., Gutarra, S., Ballestar, E., Serrano, A.L., et al. (2016). Autophagy maintains stemness by preventing senescence. Nature 529, 37–42. 10.1038/nature16187.

7. Zhang, H., Alsaleh, G., Feltham, J., Sun, Y., Napolitano, G., Riffelmacher, T., Charles, P., Frau, L., Hublitz, P., Yu, Z., et al. (2019). Polyamines Control eIF5A Hypusination, TFEB Translation, and Autophagy to Reverse B Cell Senescence. Mol. Cell 76, 110–125.e9. 10.1016/j.molcel.2019.08.005.

8. Moreno-Jiménez, E.P., Flor-García, M., Terreros-Roncal, J., Rábano, A., Cafini, F., Pallas-Bazarra, N., Ávila, J., and Llorens-Martín, M. (2019). Adult hippocampal neurogenesis is abundant in neurologically healthy subjects and drops sharply in patients with Alzheimer’s disease. Nat. Med. 25, 554–560. 10.1038/s41591-019-0375-9.

9. Baugh, L.R. (2013). To grow or not to grow: Nutritional control of development during Caenorhabditis elegans L1 Arrest at Genetics, 10.1534/genetics.113.150847 https://doi.org/10.1534/genetics.113.150847.

10. Baugh, L.R., and Sternberg, P.W. (2006). DAF-16/FOXO regulates transcription of cki-1/Cip/Kip and repression of lin-4 during C. elegans L1 arrest. Curr. Biol. 16, 780–785. 10.1016/j.cub.2006.03.021.

11. Kaplan, R.E.W., Webster, A.K., Chitrakar, R., Dent, J.A., and Baugh, L.R. (2018). Food perception without ingestion leads to metabolic changes and irreversible developmental arrest in C. elegans. BMC Biol. 16, 1–16. 10.1186/s12915-018-0579-3.

12. Roux, A.E., Langhans, K., Huynh, W., and Kenyon, C. (2016). Reversible Age-Related Phenotypes Induced during Larval Quiescence in C. Elegans. Cell Metab. 10.1016/j.cmet.2016.05.024.

13. Rodgers, J.T., King, K.Y., Brett, J.O., Cromie, M.J., Charville, G.W., Maguire, K.K., Brunson, C., Mastey, N., Liu, L., Tsai, C.-R., et al. (2014). mTORC1 controls the adaptive transition of quiescent stem cells from G0 to G(Alert). Nature 510, 393–396. 10.1038/nature13255.

14. Adhikari, D., Zheng, W., Shen, Y., Gorre, N., Hämäläinen, T., Cooney, A.J., Huhtaniemi, I., Lan, Z.-J., and Liu, K. (2010). Tsc/mTORC1 signaling in oocytes governs the quiescence and activation of primordial follicles. Hum. Mol. Genet. 19, 397–410. 10.1093/hmg/ddp483.

15. Leontieva, O. V, Demidenko, Z.N., and Blagosklonny, M. V (2014). Contact inhibition and high cell density deactivate the mammalian target of rapamycin pathway, thus suppressing the senescence program. Proc. Natl. Acad. Sci. U. S. A. 111, 8832–8837. 10.1073/pnas.1405723111.

16. Carroll, B., Nelson, G., Rabanal-Ruiz, Y., Kucheryavenko, O., Dunhill-Turner, N.A., Chesterman, C.C., Zahari, Q., Zhang, T., Conduit, S.E., Mitchell, C.A., et al. (2017). Persistent mTORC1 signaling in cell senescence results from defects in amino acid and growth factor sensing. J. Cell Biol. 216, 1949–1957. 10.1083/jcb.201610113.

17. Frakes, A.E., and Dillin, A. (2017). The UPRER: Sensor and Coordinator of Organismal Homeostasis, 10.1016/j.molcel.2017.05.031 https://doi.org/10.1016/j.molcel.2017.05.031.

18. Martínez, G., Duran-Aniotz, C., Cabral-Miranda, F., Vivar, J.P., and Hetz, C. (2017). Endoplasmic reticulum proteostasis impairment in aging. Aging Cell 16, 615–623. 10.1111/acel.12599.

19. Walter, P., and Ron, D. (2011). The unfolded protein response: from stress pathway to homeostatic regulation. Science 334, 1081–1086. 10.1126/science.1209038.

20. Hetz, C., Zhang, K., and Kaufman, R.J. (2020). Mechanisms, regulation and functions of the unfolded protein response. Nat. Rev. Mol. Cell Biol. 21, 421–438. 10.1038/s41580-020-0250-z.

21. Hetz, C., Axten, J.M., and Patterson, J.B. (2019). Pharmacological targeting of the unfolded protein response for disease intervention. Nat. Chem. Biol. 15, 764–775. 10.1038/s41589-019-0326-2.

22. Liu, Y., Shao, M., Wu, Y., Yan, C., Jiang, S., Liu, J., Dai, J., Yang, L., Li, J., Jia, W., et al. (2015). Role for the endoplasmic reticulum stress sensor IRE1α in liver regenerative responses. J. Hepatol. 10.1016/j.jhep.2014.10.022.

23. Dong, S., Wang, Q., Kao, Y.-R., Diaz, A., Tasset, I., Kaushik, S., Thiruthuvanathan, V., Zintiridou, A., Nieves, E., Dzieciatkowska, M., et al. (2021). Chaperone-mediated autophagy sustains haematopoietic stem-cell function. Nature 591, 117–123. 10.1038/s41586-020-03129-z.

24. Chen, J., Ou, Y., Li, Y., Hu, S., Shao, L.-W., and Liu, Y. (2017). Metformin extends C. elegans lifespan through lysosomal pathway. Elife 6. 10.7554/eLife.31268.

25. Smith, H.J., Lanjuin, A., Sharma, A., Prabhakar, A., Nowak, E., Stine, P.G., Sehgal, R., Stojanovski, K., Towbin, B.D., and Mair, W.B. (2023). Neuronal mTORC1 inhibition promotes longevity without suppressing anabolic growth and reproduction in C. elegans. PLOS Genet. 19, e1010938.

26. Liu, G.Y., and Sabatini, D.M. (2020). mTOR at the nexus of nutrition, growth, ageing and disease. Nat. Rev. Mol. Cell Biol. 21, 183–203. 10.1038/s41580-019-0199-y.

27. Yamamoto, H., Zhang, S., and Mizushima, N. (2023). Autophagy genes in biology and disease. Nat. Rev. Genet. 24, 382–400. 10.1038/s41576-022-00562-w.

28. Olivas, T.J., Wu, Y., Yu, S., Luan, L., Choi, P., Guinn, E.D., Nag, S., De Camilli, P. V, Gupta, K., and Melia, T.J. (2023). ATG9 vesicles comprise the seed membrane of mammalian autophagosomes. J. Cell Biol. 222. 10.1083/jcb.202208088.

29. Sawa-Makarska, J., Baumann, V., Coudevylle, N., von Bülow, S., Nogellova, V., Abert, C., Schuschnig, M., Graef, M., Hummer, G., and Martens, S. (2020). Reconstitution of autophagosome nucleation defines Atg9 vesicles as seeds for membrane formation. Science 369. 10.1126/science.aaz7714.

30. Matoba, K., Kotani, T., Tsutsumi, A., Tsuji, T., Mori, T., Noshiro, D., Sugita, Y., Nomura, N., Iwata, S., Ohsumi, Y., et al. (2020). Atg9 is a lipid scramblase that mediates autophagosomal membrane expansion. Nat. Struct. Mol. Biol. 27, 1185–1193. 10.1038/s41594-020-00518-w.

31. Ghanbarpour, A., Valverde, D.P., Melia, T.J., and Reinisch, K.M. (2021). A model for a partnership of lipid transfer proteins and scramblases in membrane expansion and organelle biogenesis. Proc. Natl. Acad. Sci. U. S. A. 118. 10.1073/pnas.2101562118.

32. Wu, F., Li, Y., Wang, F., Noda, N.N., and Zhang, H. (2012). Differential function of the two Atg4 homologues in the aggrephagy pathway in Caenorhabditis elegans. J. Biol. Chem. 287, 29457–29467. 10.1074/jbc.M112.365676.

33. Hill, S.E., Kauffman, K.J., Krout, M., Richmond, J.E., Melia, T.J., and Colón-Ramos, D.A. (2019). Maturation and Clearance of Autophagosomes in Neurons Depends on a Specific Cysteine Protease Isoform, ATG-4.2. Dev. Cell 49, 251–266.e8. 10.1016/j.devcel.2019.02.013.

34. Zhang, H., Chang, J.T., Guo, B., Hansen, M., Jia, K., Kovács, A.L., Kumsta, C., Lapierre, L.R., Legouis, R., Lin, L., et al. (2015). Guidelines for monitoring autophagy in Caenorhabditis elegans. Autophagy 11, 9–27. 10.1080/15548627.2014.1003478.

35. Tian, Y., Li, Z., Hu, W., Ren, H., Tian, E., Zhao, Y., Lu, Q., Huang, X., and Yang, P. (2010). C. elegans Screen Identifies Autophagy Genes Specific to Multicellular Organisms. 1042–1055. 10.1016/j.cell.2010.04.034.

36. Subramani, S., and Malhotra, V. (2013). Non-autophagic roles of autophagy-related proteins. EMBO Rep. 14, 143–151. 10.1038/embor.2012.220.

37. Hibshman, J.D., Leuthner, T.C., Shoben, C., Mello, D.F., Sherwood, D.R., Meyer, J.N., and Baugh, L.R. (2018). Nonselective autophagy reduces mitochondrial content during starvation in Caenorhabditis elegans. Am. J. Physiol. Cell Physiol. 315, C781–C792. 10.1152/ajpcell.00109.2018.

38. Uma Naresh, N., Kim, S., Shpilka, T., Yang, Q., Du, Y., and Haynes, C.M. (2022). Mitochondrial genome recovery by ATFS-1 is essential for development after starvation. Cell Rep. 41, 111875. 10.1016/j.celrep.2022.111875.

39. Murley, A., Lackner, L.L., Osman, C., West, M., Voeltz, G.K., Walter, P., and Nunnari, J. (2013). ER-associated mitochondrial division links the distribution of mitochondria and mitochondrial DNA in yeast. Elife. 10.7554/eLife.00422.

40. Lewis, S.C., Uchiyama, L.F., and Nunnari, J. (2016). ER-mitochondria contacts couple mtDNA synthesis with Mitochondrial division in human cells. Science (80-.). 10.1126/science.aaf5549.

41. Pickrell, A.M., and Youle, R.J. (2015). The roles of PINK1, parkin, and mitochondrial fidelity in Parkinson’s disease. Neuron 85, 257–273.

42. Wei, J., Long, L., Yang, K., Guy, C., Shrestha, S., Chen, Z., Wu, C., Vogel, P., Neale, G., Green, D.R., et al. (2016). Autophagy enforces functional integrity of regulatory T cells by coupling environmental cues and metabolic homeostasis. Nat. Immunol. 17, 277–285. 10.1038/ni.3365.

43. Bialik, S., Dasari, S.K., and Kimchi, A. (2018). Autophagy-dependent cell death–where, how and why a cell eats itself to death. J. Cell Sci. 131, jcs215152.

44. Napolitano, G., and Ballabio, A. (2016). TFEB at a glance. J. Cell Sci. 10.1242/jcs.146365.

45. Lapierre, L.R., De Magalhaes Filho, C.D., McQuary, P.R., Chu, C.-C., Visvikis, O., Chang, J.T., Gelino, S., Ong, B., Davis, A.E., Irazoqui, J.E., et al. (2013). The TFEB orthologue HLH-30 regulates autophagy and modulates longevity in Caenorhabditis elegans. Nat. Commun. 4, 2267. 10.1038/ncomms3267.

46. Springhorn, A., and Hoppe, T. (2019). Western blot analysis of the autophagosomal membrane protein LGG-1/LC3 in Caenorhabditis elegans. Methods Enzymol. 619, 319–336. 10.1016/bs.mie.2018.12.034.

47. Lin, L., Yang, P., Huang, X., Zhang, H., Lu, Q., and Zhang, H. (2013). The scaffold protein EPG-7 links cargo-receptor complexes with the autophagic assembly machinery. J. Cell Biol. 201, 113–129. 10.1083/jcb.201209098.

48. Jia, J., Abudu, Y.P., Claude-Taupin, A., Gu, Y., Kumar, S., Choi, S.W., Peters, R., Mudd, M.H., Allers, L., Salemi, M., et al. (2018). Galectins Control mTOR in Response to Endomembrane Damage. Mol. Cell 70, 120–135.e8. 10.1016/j.molcel.2018.03.009.

49. Li, Y., Chen, B., Zou, W., Wang, X., Wu, Y., Zhao, D., Sun, Y., Liu, Y., Chen, L., Miao, L., et al. (2016). The lysosomal membrane protein SCAV-3 maintains lysosome integrity and adult longevity. J. Cell Biol. 215, 167–185. 10.1083/jcb.201602090.

50. Flavin, W.P., Bousset, L., Green, Z.C., Chu, Y., Skarpathiotis, S., Chaney, M.J., Kordower, J.H., Melki, R., and Campbell, E.M. (2017). Endocytic vesicle rupture is a conserved mechanism of cellular invasion by amyloid proteins. Acta Neuropathol. 134, 629–653. 10.1007/s00401-017-1722-x.

51. Freeman, D., Cedillos, R., Choyke, S., Lukic, Z., McGuire, K., Marvin, S., Burrage, A.M., Sudholt, S., Rana, A., O’Connor, C., et al. (2013). Alpha-synuclein induces lysosomal rupture and cathepsin dependent reactive oxygen species following endocytosis. PLoS One 8, e62143. 10.1371/journal.pone.0062143.

52. D’Astolfo, D.S., Pagliero, R.J., Pras, A., Karthaus, W.R., Clevers, H., Prasad, V., Lebbink, R.J., Rehmann, H., and Geijsen, N. (2015). Efficient Intracellular Delivery of Native Proteins. Cell 161, 674– 690. 10.1016/j.cell.2015.03.028.

53. Todaro, G.J., and Green, H. (1963). Quantitative studies of the growth of mouse embryo cells in culture and their development into established lines. J. Cell Biol. 17, 299–313. 10.1083/jcb.17.2.299.

54. Reznikoff, C.A., Brankow, D.W., and Heidelberger, C. (1973). Establishment and characterization of a cloned line of C3H mouse embryo cells sensitive to postconfluence inhibition of division. Cancer Res. 33, 3231–3238.

55. Dunn, K.C., Aotaki-Keen, A.E., Putkey, F.R., and Hjelmeland, L.M. (1996). ARPE-19, A Human Retinal Pigment Epithelial Cell Line with Differentiated Properties. Exp. Eye Res. 62, 155–170. 10.1006/exer.1996.0020.

56. Harnoss, J.M., Le Thomas, A., Shemorry, A., Marsters, S.A., Lawrence, D.A., Lu, M., Chen, Y.-C.A., Qing, J., Totpal, K., Kan, D., et al. (2019). Disruption of IRE1α through its kinase domain attenuates multiple myeloma. Proc. Natl. Acad. Sci. U. S. A. 116, 16420–16429. 10.1073/pnas.1906999116.

57. Harrington, P.E., Biswas, K., Malwitz, D., Tasker, A.S., Mohr, C., Andrews, K.L., Dellamaggiore, K., Kendall, R., Beckmann, H., Jaeckel, P., et al. (2015). Unfolded Protein Response in Cancer: IRE1α Inhibition by Selective Kinase Ligands Does Not Impair Tumor Cell Viability. ACS Med. Chem. Lett. 6, 68–72. 10.1021/ml500315b.

58. Hansen, M., Rubinsztein, D.C., and Walker, D.W. (2018). Autophagy as a promoter of longevity: insights from model organisms. Nat. Rev. Mol. Cell Biol. 19, 579–593. 10.1038/s41580-018-0033-y.

59. Chang, J.T., Kumsta, C., Hellman, A.B., Adams, L.M., and Hansen, M. (2017). Spatiotemporal regulation of autophagy during Caenorhabditis elegans aging. Elife 6. 10.7554/eLife.18459.

60. Fujimaki, K., Li, R., Chen, H., Croce, K. Della, Zhang, H.H., Xing, J., Bai, F., and Yao, G. (2019). Graded regulation of cellular quiescence depth between proliferation and senescence by a lysosomal dimmer switch. Proc. Natl. Acad. Sci. U. S. A. 116, 22624–22634. 10.1073/pnas.1915905116.

61. Young, A.R.J., Narita, M., Ferreira, M., Kirschner, K., Sadaie, M., Darot, J.F.J., Tavaré, S., Arakawa, S., Shimizu, S., Watt, F.M., et al. (2009). Autophagy mediates the mitotic senescence transition. Genes Dev. 23, 798–803. 10.1101/gad.519709.

62. Johmura, Y., Yamanaka, T., Omori, S., Wang, T.-W., Sugiura, Y., Matsumoto, M., Suzuki, N., Kumamoto, S., Yamaguchi, K., Hatakeyama, S., et al. (2021). Senolysis by glutaminolysis inhibition ameliorates various age-associated disorders. Science (80-.). 371, 265 LP – 270. 10.1126/science.abb5916.

63. Eapen, V. V, Swarup, S., Hoyer, M.J., Paulo, J.A., and Harper, J.W. (2021). Quantitative proteomics reveals the selectivity of ubiquitin-binding autophagy receptors in the turnover of damaged lysosomes by lysophagy. Elife 10. 10.7554/eLife.72328.

64. Zoncu, R., and Perera, R.M. (2022). Built to last: lysosome remodeling and repair in health and disease. Trends Cell Biol. 32, 597–610. 10.1016/j.tcb.2021.12.009.

65. Tang, A.H., and Rando, T.A. (2014). Induction of autophagy supports the bioenergetic demands of quiescent muscle stem cell activation. EMBO J. 33, 2782–2797. 10.15252/embj.201488278.

66. Nakamura, S., Shigeyama, S., Minami, S., Shima, T., Akayama, S., Matsuda, T., Esposito, A., Napolitano, G., Kuma, A., Namba-Hamano, T., et al. (2020). LC3 lipidation is essential for TFEB activation during the lysosomal damage response to kidney injury. Nat. Cell Biol. 22, 1252–1263. 10.1038/s41556-020-00583-9.

67. Borsa, M., Obba, S., Richter, F.C., Zhang, H., Riffelmacher, T., Carrelha, J., Alsaleh, G., Jacobsen, S.E.W., and Simon, A.K. (2024). Autophagy preserves hematopoietic stem cells by restraining MTORC1-mediated cellular anabolism. Autophagy 20, 45–57. 10.1080/15548627.2023.2247310.

68. Mainz, L., Sarhan, M.A.F.E., Roth, S., Sauer, U., Kalogirou, C., Eckstein, M., Gerhard-Hartmann, E., Seibert, H.-D., Voelker, H.-U., Geppert, C., et al. (2022). Acute systemic knockdown of Atg7 is lethal and causes pancreatic destruction in shRNA transgenic mice. Autophagy, 1–14. 10.1080/15548627.2022.2052588.

69. Li, T.Y., Gao, A.W., Li, X., Liu, Y.J., Arey, R.N., Morales, K., Lalou, A., Wang, Q., Lima, T., and Auwerx, J. (2022). A lysosomal surveillance response (LySR) that reduces proteotoxicity and extends healthspan. bioRxiv, 2022.06.13.495962. 10.1101/2022.06.13.495962.

70. Song, J., Ni, Q., Sun, J., Xie, J., Liu, J., Ning, G., Wang, W., and Wang, Q. (2022). Aging Impairs Adaptive Unfolded Protein Response and Drives Beta Cell Dedifferentiation in Humans. J. Clin. Endocrinol. Metab. 107, 3231–3241. 10.1210/clinem/dgac535.

71. Taylor, R.C., and Dillin, A. (2013). XBP-1 is a cell-nonautonomous regulator of stress resistance and longevity. Cell 153, 1435–1447. 10.1016/j.cell.2013.05.042.

72. Sabath, N., Levy-Adam, F., Younis, A., Rozales, K., Meller, A., Hadar, S., Soueid-Baumgarten, S., and Shalgi, R. (2020). Cellular proteostasis decline in human senescence. Proc. Natl. Acad. Sci. U. S. A. 117, 31902–31913. 10.1073/pnas.2018138117.

73. Paz Gavilán, M., Pintado, C., Gavilán, E., Jiménez, S., Ríos, R.M., Vitorica, J., Castaño, A., and Ruano, D. (2009). Dysfunction of the unfolded protein response increases neurodegeneration in aged rat hippocampus following proteasome inhibition. Aging Cell 8, 654–665. 10.1111/j.1474-9726.2009.00519.x.

74. Lu, F., Leach, L.L., and Gross, J.M. (2022). mTOR activity is essential for retinal pigment epithelium regeneration in zebrafish. PLoS Genet. 18, e1009628. 10.1371/journal.pgen.1009628.

75. Curnock, R., Yalci, K., Palmfeldt, J., Jäättelä, M., Liu, B., and Carroll, B. (2023). TFEB-dependent lysosome biogenesis is required for senescence. EMBO J. 42, e111241. 10.15252/embj.2022111241.

76. Chen, J.J., Nathaniel, D.L., Raghavan, P., Nelson, M., Tian, R., Tse, E., Hong, J.Y., See, S.K., Mok, S.-A., Hein, M.Y., et al. (2019). Compromised function of the ESCRT pathway promotes endolysosomal escape of tau seeds and propagation of tau aggregation. J. Biol. Chem. 294, 18952–18966. 10.1074/jbc.RA119.009432.

77. Vaquer-Alicea, J., and Diamond, M.I. (2019). Propagation of Protein Aggregation in Neurodegenerative Diseases. Annu. Rev. Biochem. 88, 785–810. 10.1146/annurev-biochem-061516-045049.

78. Calafate, S., Flavin, W., Verstreken, P., and Moechars, D. (2016). Loss of Bin1 Promotes the Propagation of Tau Pathology. Cell Rep. 17, 931–940. 10.1016/j.celrep.2016.09.063.

79. Rose, K., Jepson, T., Shukla, S., Maya-Romero, A., Kampmann, M., Xu, K., and Hurley, J.H. (2024). Tau fibrils induce nanoscale membrane damage and nucleate cytosolic tau at lysosomes. Proc. Natl. Acad. Sci. U. S. A. 121, e2315690121. 10.1073/pnas.2315690121.

80. Burbidge, K., Rademacher, D.J., Mattick, J., Zack, S., Grillini, A., Bousset, L., Kwon, O., Kubicki, K., Simon, A., Melki, R., et al. (2022). LGALS3 (galectin 3) mediates an unconventional secretion of SNCA/α-synuclein in response to lysosomal membrane damage by the autophagic-lysosomal pathway in human midbrain dopamine neurons. Autophagy 18, 1020–1048. 10.1080/15548627.2021.1967615.

81. Silva-García, C.G., Lanjuin, A., Heintz, C., Dutta, S., Clark, N.M., and Mair, W.B. (2019). Single-Copy Knock-In Loci for Defined Gene Expression in Caenorhabditis elegans. G3 (Bethesda). 9, 2195–2198. 10.1534/g3.119.400314.

82. Lehrbach, N.J., Ji, F., and Sadreyev, R. (2017). Next-Generation Sequencing for Identification of EMS-Induced Mutations in Caenorhabditis elegans. Curr. Protoc. Mol. Biol. 117, 7.29.1–7.29.12. 10.1002/cpmb.27.

83. Frankino, P.A., Siddiqi, T.F., Bolas, T., Bar-Ziv, R., Gildea, H.K., Zhang, H., Higuchi-Sanabria, R., and Dillin, A. (2022). SKN-1 regulates stress resistance downstream of amino catabolism pathways. iScience 25, 104571. 10.1016/j.isci.2022.104571.

84. The Galaxy platform for accessible, reproducible and collaborative biomedical analyses: 2022 update (2022). Nucleic Acids Res. 50, W345–W351. 10.1093/nar/gkac247.

85. Moos, K., Seifert, M., Baumeister, R., Maier, W. MiModD. https://mimodd.readthedocs.io/en/latest/.

86. Cantor, J.R., Abu-Remaileh, M., Kanarek, N., Freinkman, E., Gao, X., Louissaint, A., Lewis, C.A., and Sabatini, D.M. (2017). Physiologic Medium Rewires Cellular Metabolism and Reveals Uric Acid as an Endogenous Inhibitor of UMP Synthase. Cell 169, 258–272.e17. 10.1016/j.cell.2017.03.023.

87. Chen, B., Gilbert, L.A., Cimini, B.A., Schnitzbauer, J., Zhang, W., Li, G.-W., Park, J., Blackburn, E.H., Weissman, J.S., and Qi, L.S. (2013). Dynamic imaging of genomic loci in living human cells by an optimized CRISPR/Cas system. Cell 155, 1479–1491.

88. Sanjana, N.E., Shalem, O., and Zhang, F. (2014). Improved vectors and genome-wide libraries for CRISPR screening., 10.1038/nmeth.3047 https://doi.org/10.1038/nmeth.3047.

89. Replogle, J.M., Bonnar, J.L., Pogson, A.N., Liem, C.R., Maier, N.K., Ding, Y., Russell, B.J., Wang, X., Leng, K., Guna, A., et al. (2022). Maximizing CRISPRi efficacy and accessibility with dual-sgRNA libraries and optimal effectors. Elife 11, e81856. 10.7554/eLife.81856.

