## Supplementary figures and images for "Quiescent cell re-entry is limited by macroautophagy-induced lysosomal damage"

### Supplemental Figures

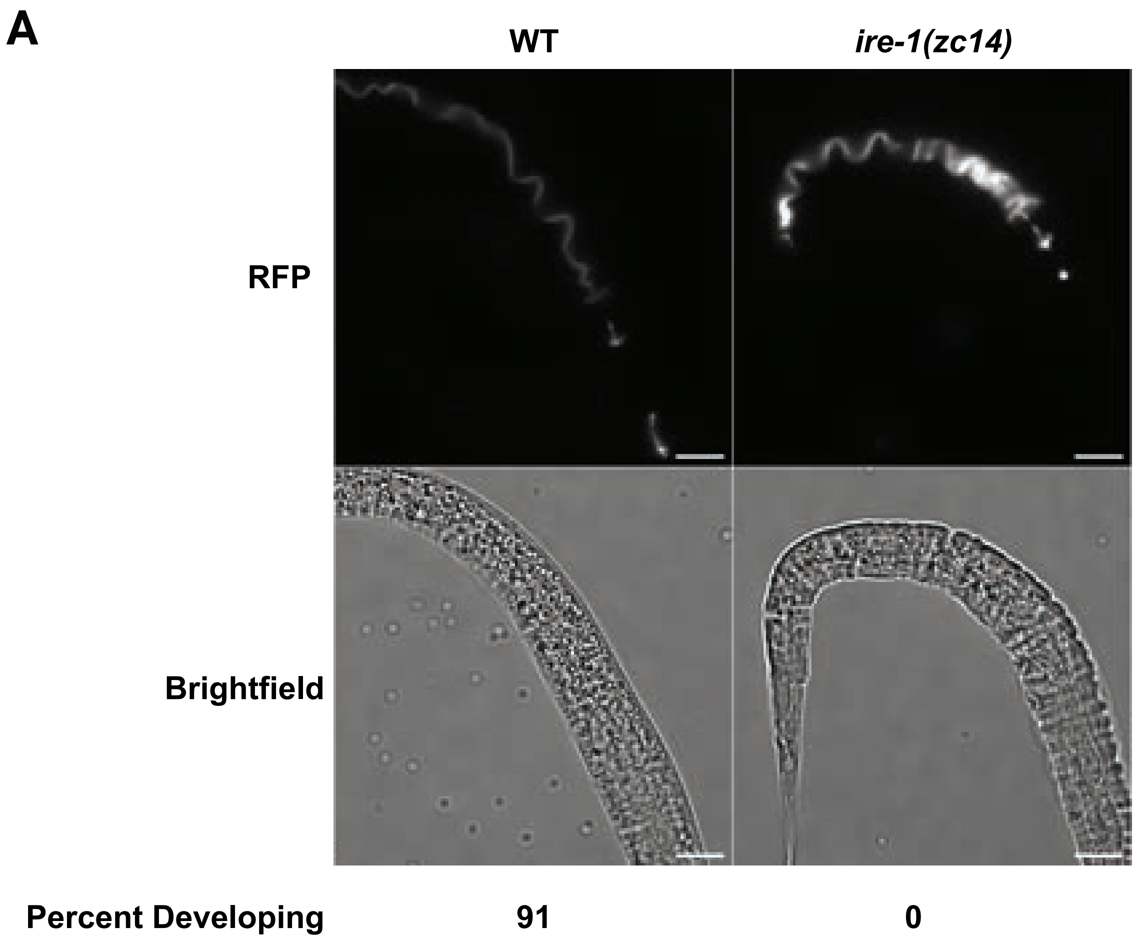

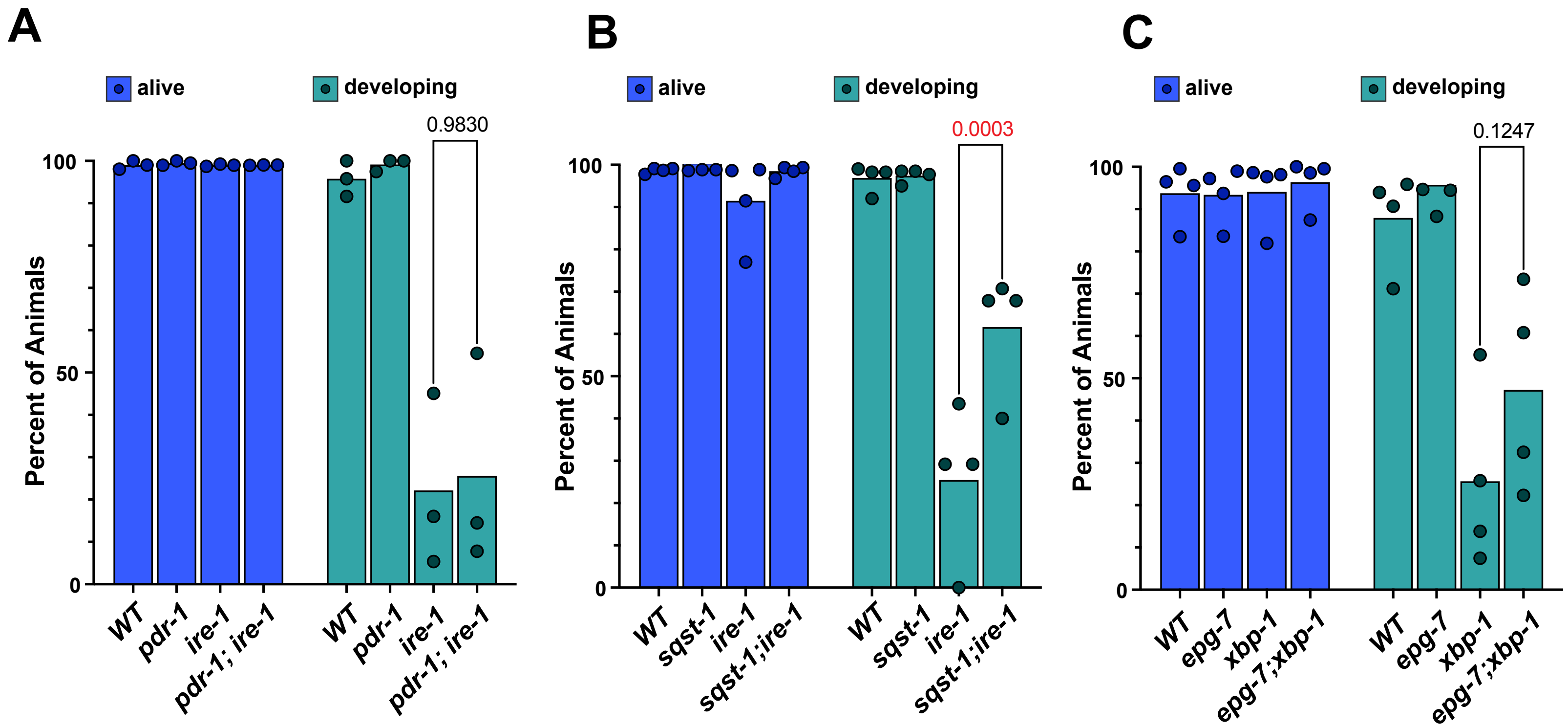

#### Figure S3

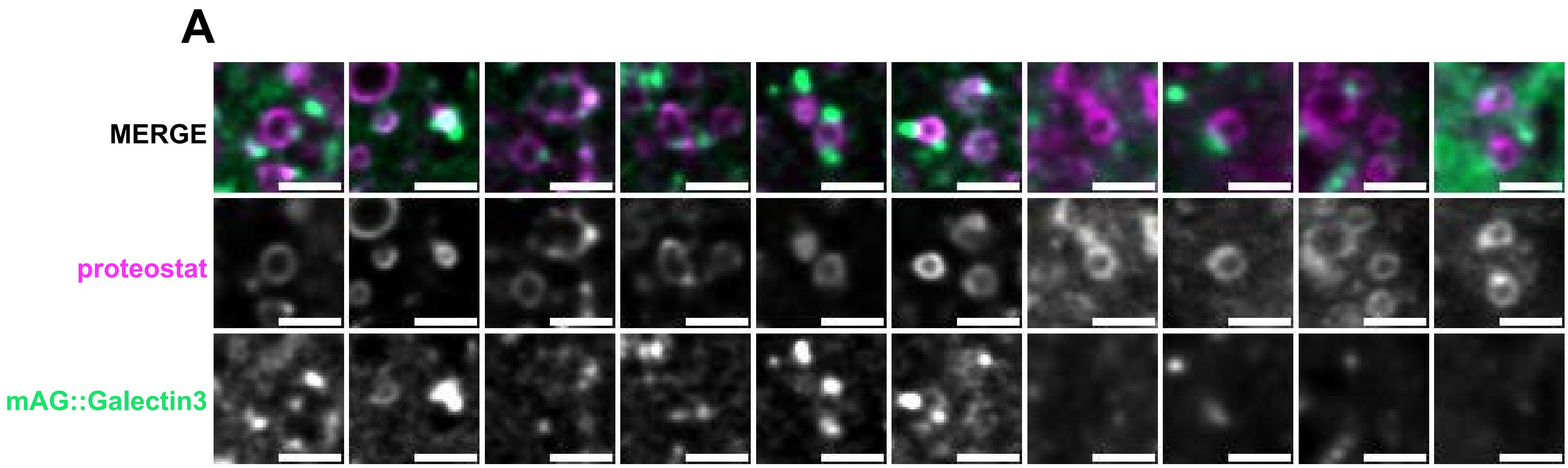

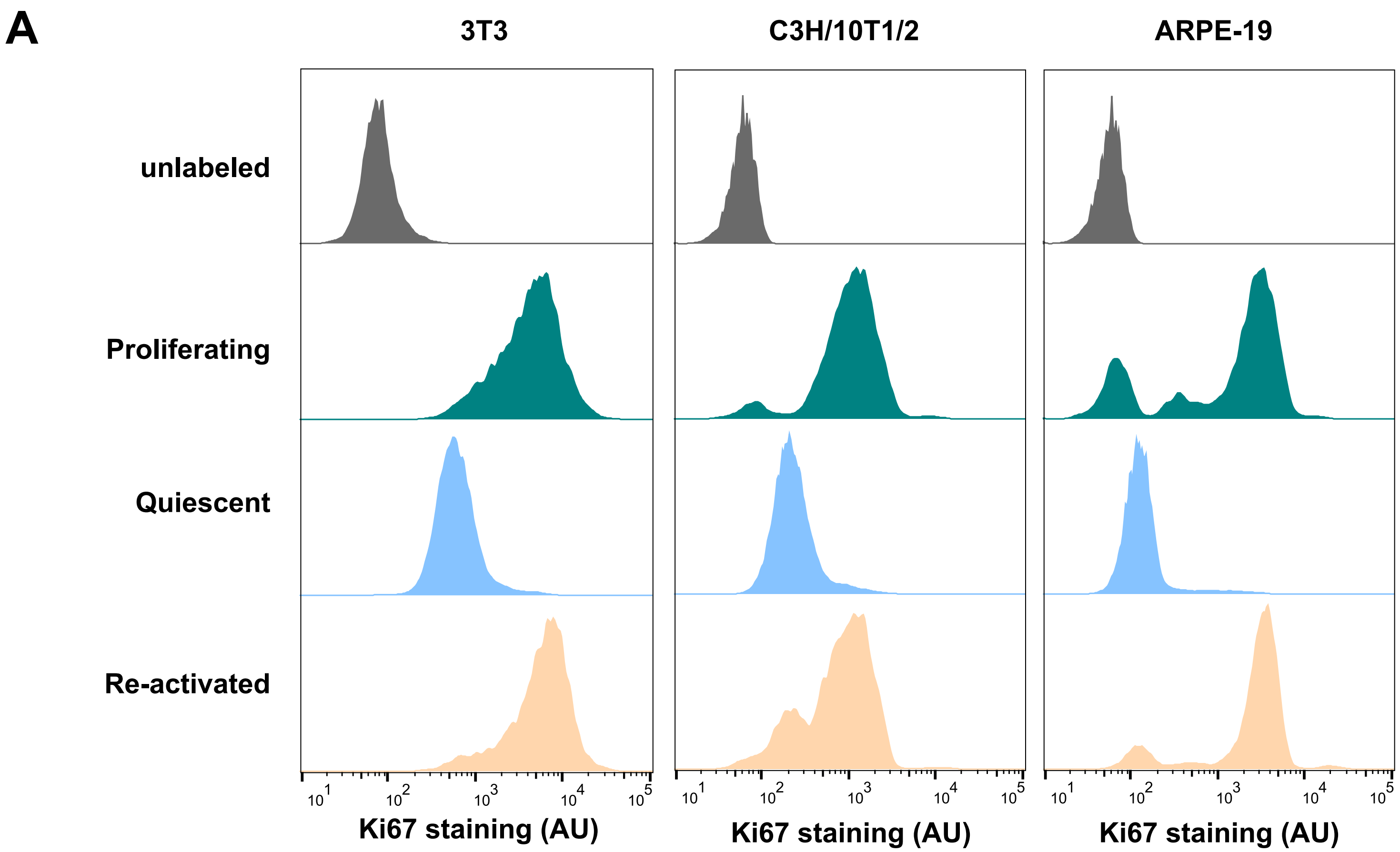

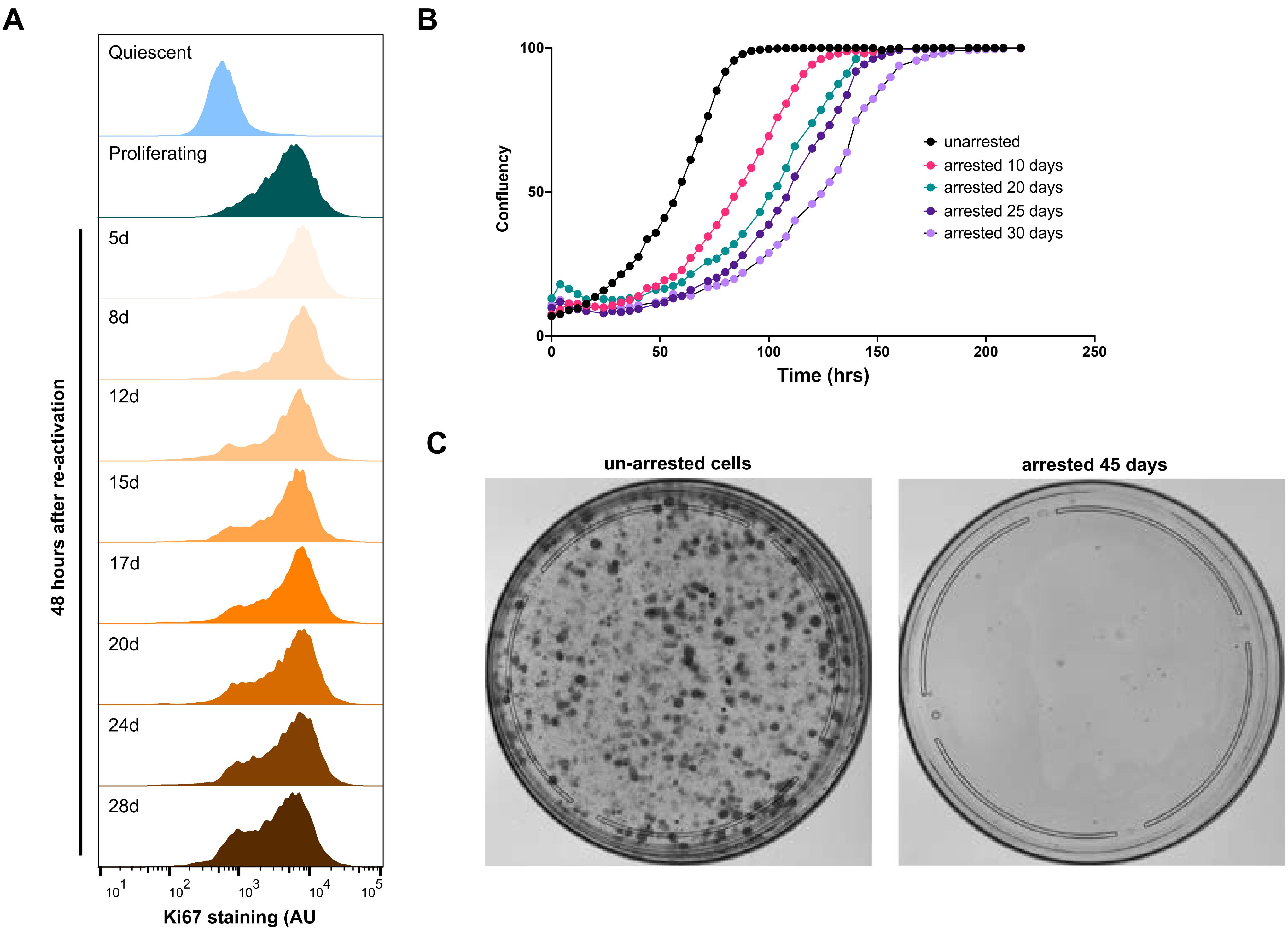

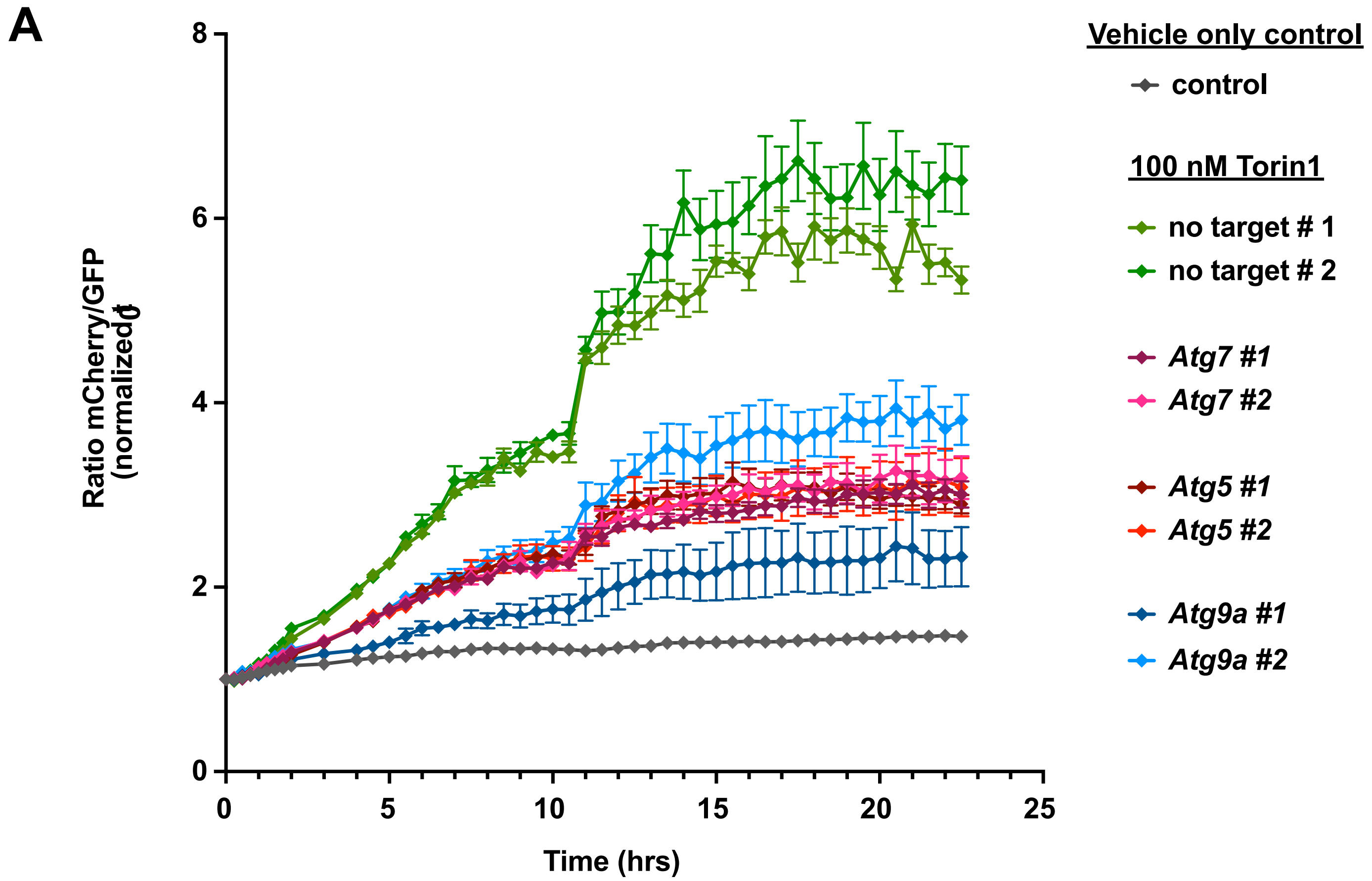
